# Characterization of DNA methylation reader proteins of *Arabidopsis thaliana*

**DOI:** 10.1101/2023.12.05.570080

**Authors:** Jonathan Cahn, James P. B. Lloyd, Ino D. Karemaker, Pascal W.T.C. Jansen, Jahnvi Pflueger, Owen Duncan, Jakob Petereit, Ozren Bogdanovic, A. Harvey Millar, Michiel Vermeulen, Ryan Lister

## Abstract

In plants, cytosine DNA methylation (mC) is largely associated with transcriptional repression of transposable elements, but it can also be found in the body of expressed genes, referred to as gene body methylation (GbM). GbM is correlated with ubiquitously expressed genes, however its function, or absence thereof, is highly debated. The different output that mC can have raises questions as to how it is interpreted - or read - differently in these sequence and genomic contexts. To screen for potential mC binding proteins, we performed an unbiased DNA affinity pull-down assay combined with quantitative mass spectrometry using methylated DNA probes for each DNA sequence context. All mC readers known to date were found to preferentially bind to the methylated probes, along with a range of new mC binding protein candidates. Functional characterization of these mC readers, focused on the MBD and SUVH families, was undertaken by ChIP-seq mapping of genome-wide binding sites, their protein interactors, and the impact of high-order mutations on transcriptomic and epigenomic profiles. Together, this highlighted specific context preferences for these proteins, and in particular the ability of MBD2 to bind specifically to GbM. This comprehensive analysis of Arabidopsis mC readers emphasizes the complexity and interconnectivity between DNA methylation and chromatin remodelling processes in plants.

## Introduction

Epigenomic modifications constitute additional regulatory layers that can alter and/or record transcriptional activity of their underlying genetic sequences (Bird 2007). One such modification is cytosine DNA methylation (mC), the covalent bond of a methyl group (-CH_3_) to the 5^th^ carbon of cytosine bases. In plants, mC can occur in three DNA sequence contexts: CG, CHG, and CHH (H = A, C, or T). Methylation in each sequence context shows distinct distribution patterns along the genome depending on underlying sequence features and elements (Lloyd and Lister 2022), and is regulated by different enzymes, with DNA methyltransferases responsible for mC deposition (‘writers’), and DNA glycosylases catalyzing DNA demethylation (‘erasers’). Investigation of DNA methyltransferases in Arabidopsis has revealed sequence context-specific activity for these enzymes: MET1 is responsible for maintaining mCG, being recruited to hemi-methylated sites after replication (Finnegan et al. 1996; Ronemus et al. 1996; Shook and Richards 2014); CMT3 and CMT2 maintain mCHG and mCHH, respectively, via feed-forward loops with the histone post-transcriptional modification H3K9me2 (Lindroth et al. 2001; Jackson et al. 2002; Du et al. 2012; Zemach et al. 2013; Stroud et al. 2014); and DRM2 is required for *de novo* DNA methylation in all three contexts through the RNA-directed DNA methylation (RdDM) pathway (Cao and Jacobsen 2002; Matzke and Mosher 2014; Lloyd and Lister 2022). DNA demethylation appears to exhibit less sequence context specificity, where DME is involved in all demethylation events in the companion cells of the plant gametes but primarily in the CG context, and ROS1, DML2, and DML3 are partially redundant in vegetative tissues for all mC contexts (Gong et al. 2002; Gehring et al. 2006; Penterman et al. 2007; Hsieh et al. 2009; Ibarra et al. 2012).

In Arabidopsis, heterochromatin is marked by high DNA methylation in all three contexts and is associated with transcriptional silencing of transposable elements (TEs) (Cokus et al. 2008; Lister et al. 2008; Lloyd and Lister 2022). In addition to an important role in silencing TEs in plants, notably in germline cells, mC is also involved in other aspects of plant development, such as imprinting by regulating MEDEA, a histone methyltransferase of the polycomb repressive complex 2 (Satyaki and Gehring 2017), in recombination patterns during meiosis, where mCHG limits heterochromatin rearrangements (Underwood et al. 2018), or in controlling alternative splicing in male sex cells of meiosis factor *MPS1* (Walker et al. 2018). Many genes (10-20%)(Zhang et al. 2020) in euchromatic arms also harbor DNA methylation but only in the CG context, located in the gene body with a modest bias towards the 3’ end, called gene body methylation (GbM) (Zhang et al. 2006; Cokus et al. 2008; Lister et al. 2008). In contrast with heterochromatic mC, GbM is associated with ubiquitously and moderately expressed genes in plants (Zhang et al. 2006; Takuno and Gaut 2012; Coleman-Derr and Zilberman 2012). Despite being present in almost all land plants, whether GbM has a molecular function has been highly debated since some plant species do not harbor GbM (Niederhuth et al. 2016; Bewick et al. 2016; Zilberman 2017). A current hypothesis proposes that GbM is a co-product of stable mC maintenance, and the lack of a described mechanism that specifically regulates GbM to date would currently support this as the most parsimonious explanation (Bewick et al. 2016; Bewick and Schmitz 2017; Bewick et al. 2017; Wendte et al. 2019).

An important factor in understanding the molecular roles of mC in different sequence and chromatin contexts is the discovery and characterisation of the proteins that can bind to DNA methylation (mC ‘readers’). Two protein domains were identified to confer the ability to bind to DNA methylation: Methyl-Binding Domain (MBD), first identified in Methyl-CpG binding protein 2 (MeCP2) (Nan et al. 1993), and the SET and RING-Associated (SRA) domain (Baumbusch et al. 2001; Johnson et al. 2007). By sequence homology with these founder proteins, three families were identified in Arabidopsis, two containing a SET-domain - three Variant In Methylation (VIM) proteins and ten Suppression of Variegation 3-9 Homolog (SUVH) proteins - and thirteen members in the MBD family (Baumbusch et al. 2001; Springer et al. 2003; Zemach and Grafi 2003; Berg et al. 2003; Springer and Kaeppler 2005). The VIM proteins can bind to mCG and mCHG *in vitro* but preferentially bind to hemi-methylated DNA in the CG sequence context (Woo et al. 2007, 2008), and are responsible for recruiting MET1 for mCG maintenance after replication (Stroud et al. 2013; Shook and Richards 2014). Several members of the SUVH family have been well characterized by their involvement in mC deposition: SUVH4/KYP, SUVH5, and SUVH6 are able to bind to mC in all contexts, but preferentially bind mCHG and mCHH (Johnson et al. 2007; Rajakumara et al. 2011; Du et al. 2014). They contain an active histone methyltransferase domain that catalyzes H3K9me2, thus forming a feed-forward loop with CMT3 and CMT2 that bind to H3K9me2 and in turn deposit mCHG and mCHH, respectively (Jackson et al. 2002; Ebbs et al. 2005; Ebbs and Bender 2006; Zemach et al. 2013; Stroud et al. 2013, 2014). SUVH2 and SUVH9 bind all methylated contexts but preferentially to mCG and mCHH, respectively (Johnson et al. 2008). They are partially redundant in the RdDM pathway, recruiting PolV for the generation of long non-coding RNA that - in concert with other factors - recruit DRM2 for *de novo* DNA methylation (Kuhlmann and Mette 2012; Johnson et al. 2014; Liu et al. 2014). SUVH1 and SUVH3 have also been identified as mC readers, and were proposed to have a role in promoting transcription of genes with hypermethylated promoters (Harris et al. 2018; Li et al. 2016; Zhao et al. 2019).

The functions and binding abilities of MBD proteins are less well understood. Like their animal counterparts, MBD proteins seem to preferentially bind to mCG, despite some reports of their binding to non-CG context methylation too (Zemach and Grafi 2003; Ito et al. 2003; Scebba et al. 2003). MBD5, MBD6, and MBD7 are the most studied members (Li et al. 2017; Ichino et al. 2021a, 2022), and have been shown to be enriched at chromocenters by GFP-fusion microscopy analysis, a localization that is dependent on the chromatin remodeler DDM1 (Zemach et al. 2005). They can interact with each other and with alpha-crystallin domain (ACD) proteins, but their molecular function remains unclear due to conflicting results regarding their involvement in chromatin remodeling or DNA demethylation (Zemach et al. 2008; Li et al. 2015, 2017). MBD8, MBD9, MBD10, and MBD11 are involved in diverse developmental processes, but based on biochemical characterisation, they are unlikely to bind to mC (Berg et al. 2003; Peng et al. 2006; Preuss et al. 2008; Yaish et al. 2009; Stangeland et al. 2009). It is unclear whether MBD1, MBD2, and MBD4 can bind to DNA methylation and their molecular functions are still unknown (Zemach and Grafi 2007), but they have recently been found in complex with histone deacetylases and may have a role in controlling flowering timing (Zhou et al. 2021).

The mC-binding ability of MBD proteins had been exclusively investigated by electrophoretic mobility shift assay, leading to some conflicting results (Zemach and Grafi 2003; Ito et al. 2003; Scebba et al. 2003). A different approach was taken more recently, by performing a DNA-affinity pull-down followed by mass spectrometry analysis (Harris et al. 2018). They used probes containing endogenous sequences of highly methylated promoters, which might have biased the discovery of proteins binding to these specific loci. In this study, we performed a DNA-affinity pull-down experiment followed by mass spectrometry using DNA probes reflecting context-specific DNA methylation states in the Arabidopsis genome, resulting in the identification of all known mC readers as well as a variety of new candidate DNA methylation binding proteins. We then focused on the characterization of their genome-wide binding pattern *in vivo*, and of high-order mutant plants lacking these proteins to interrogate the direct role of DNA methylation in Arabidopsis.

## Results

### Identification of Arabidopsis mC readers

To identify potential mC readers in Arabidopsis, we performed a DNA pull-down affinity assay of nuclear proteins using methylated DNA oligonucleotides. For each methylated sequence context (CG, CHG, CHH), DNA probes were designed to maximize the relevance of the binding proteins by searching for a representative 5 bp motif frequently found in methylated sequences of Arabidopsis genome (Supp Fig 1; see Materials and Methods). After combining probes with Arabidopsis nuclear extracts and performing affinity enrichment, mass spectrometry analysis of the pulled down proteins using a statistical analysis that compares protein enrichment for methylated and unmethylated probes of each context identified 36 proteins enriched in at least one methylated cytosine context (Fig 1; Supp Table 1). Among these proteins, all but one known mC readers were found to be enriched in binding to the methylated probes, and often in the contexts relevant to their functions (Supp Table 1). For example, the three VIM proteins were found enriched in symmetrical contexts, SUVH2 and SUVH5 were enriched in all contexts, SUVH4 (KYP) and SUVH6 were enriched in mCHG, and MBD5 and MBD6 were enriched only in mCG. Only MBD7 was excluded from our final list since it was recovered in only two out of three mCG probes. We also found expected proteins enriched for binding to unmethylated probes (Supp Table 1). For example, 13 WRKY transcription factors were enriched in the CHH probe (Fig 1b; Supp Table 2), which have previously been found to prefer binding to unmethylated DNA by DAP-seq (O’Malley et al. 2016). The CHH probe sequence contains their known target site sequence (TGAC), likely underlying their binding to this probe (O’Malley et al. 2016). This screen also identified new mC reader candidates, most notably MBD1, MBD2, and MBD4, which while related to other mC readers, have had mixed results regarding their affinity for methylated DNA in prior studies (Zemach and Grafi 2003; Scebba et al. 2003; Ito et al. 2003).

**Figure 1:**
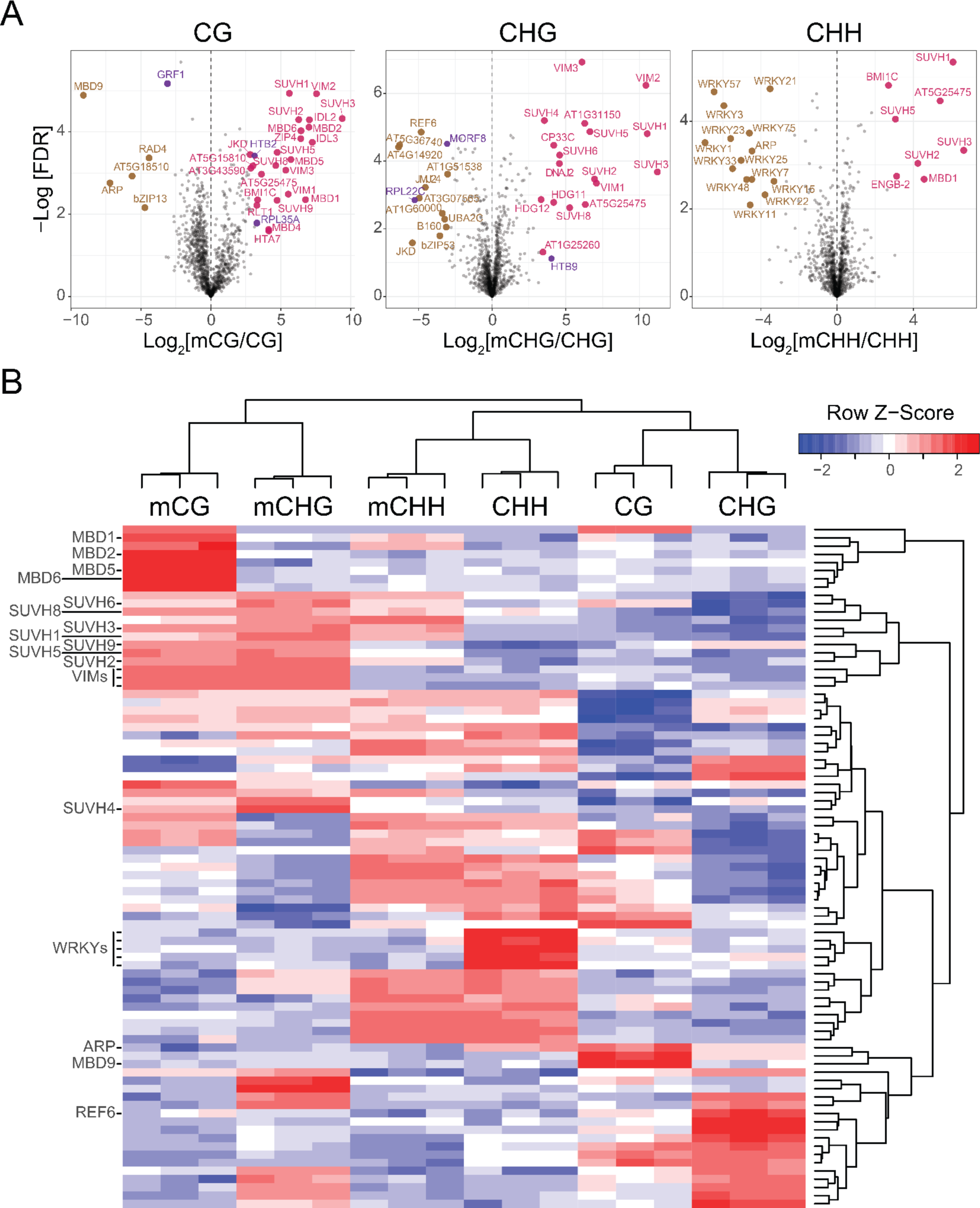
**Identification of mC reader proteins in Arabidopsis from affinity pull-downs.** A) Volcano plots of the proteins statistically enriched for binding to methylated probes (pink) or unmethylated probes (brown) in each context (CG, CHG, CHH), as well as statistically enriched proteins likely to be contaminants (purple). B) Hierarchical clustering of the proteins (rows) statistically significantly enriched in at least one set of probes (columns, each representing a replicate). The names of proteins of interest were highlighted, the rest of the 83 proteins identified can be found in Supplementary Table 2.

### Binding preferences of mC readers

To investigate the genomic binding sites of seven selected mC readers (MBD1, MBD2, MBD4, MBD5, MBD6, SUVH1, SUVH3), mutant lines of each reader (Supp Table 3) were isolated and then complemented with an epitope (2xStrep-HA-6xHis) tagged version of the corresponding protein driven by its predicted endogenous promoter. ChIP-seq against the HA epitope was performed on two independent insertion lines in T2 populations, either close to endogenous levels or overexpressed (Supp Table 3). Initial ChIP-seq peak calling analysis showed similar profiles in each replicate (Supp Fig 2). However, the line with higher protein expression had higher enrichment signals (i.e. more peaks called) and merging the replicates enabled the identification of additional peaks that would have been missed if independent replicates were used (Supp Fig 2). Thus, merged samples were used for further analyses. Peak analysis of merged replicates showed specific genome binding profiles for each protein. MBD5, MBD6, SUVH1, and SUVH3 exhibited preferential localization to heterochromatic peri-centromeric regions, whereas the other MBD proteins studied were mostly localized along euchromatic chromosome arms (Fig 2B). MBD1, MBD2, and MBD4 were mostly associated with genes, whereas MBD5, MBD6, SUVH1, and SUVH3 were primarily associated with TEs. The regions bound by MBD5, MBD6, SUVH1, and SUVH3 were highly methylated in all sequence contexts, whereas MBD2 peaks were mostly methylated in the CG context, and MBD1 peaks were surprisingly lowly methylated (Fig 2C). Sequence motif analysis (Supp Fig 3), demonstrated that, unsurprisingly, no clear motif was found for any of the candidates, indicating that their ability to bind DNA was not dependent on the genetic sequence but rather on the epigenomic state of the genomic region (Supp Fig 3). However, the most strongly enriched motifs for MBD5 and MBD6 harbor a central CG dinucleotide, which would support their preference for methylation in the CG context. The preferential binding of these proteins was also assessed *in vitro* by DNA affinity pull-down sequencing (Bartlett et al. 2017). We incubated recombinant mC reader proteins with WT Arabidopsis fragmented gDNA libraries before (DAP-seq) and after PCR amplification (ampDAP-seq), where the amplification depletes the library of DNA methylation. MBD5 and MBD6 showed sequence enrichment, but only for DAP-seq, where DNA methylation was present, and with binding site profiles that closely match those identified by ChIP-seq (Fig 2D). These results support the hypothesis that MBD5 and MBD6 binding abilities are largely or solely dictated by DNA methylation in the CG context, whereas the other mC readers tested might require other cofactors, such as protein complexes to interact with or specific epigenomic states.

**Figure 2:**
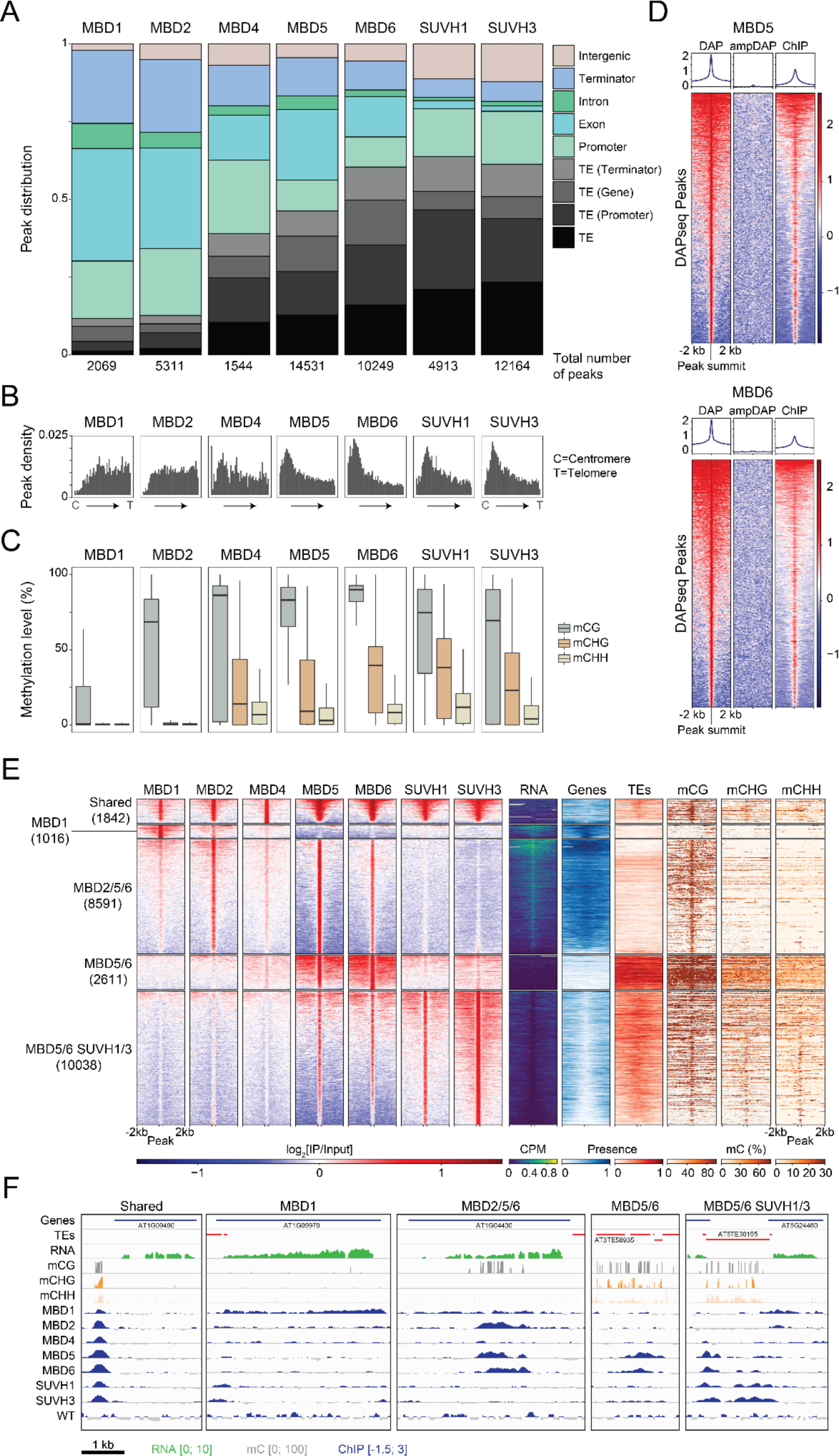
**The diverse binding profiles of mC readersacross the Arabidopsis genome.** A) Distribution of ChIP-seq peaks in genomic features. Total numbers of peaks for each sample are indicated at the top. Original annotations for TEs are indicated in parenthesis. Promoters include [-1kb; +100bp] around the transcription start site; Terminators include [-100bp; +1kb] around the transcription termination site; Genes include both exons and introns. B) Density of ChIP-seq peak positions along the Arabidopsis chromosomes, relative to their distance from the centromere (C, on the left) to the telomere of the chromosome arm they are on (T, on the right). C) Average DNA methylation levels in WT Arabidopsis seedlings under protein binding sites. Boxplots show the mean and quartiles of all peaks for each sample. For each context, only peaks containing at least 3 cytosines and a sequencing coverage ≥3 (average on all cytosines in the peak) are presented. D) DNA Affinity Purification Sequencing for MBD5 (top) and MBD6 (bottom) recombinant proteins on methylated (DAP) and unmethylated (ampDAP) genomic DNA libraries. Heatmaps show enrichment levels at all DAP-seq peaks, and the correlation with *in vivo* binding assessed by ChIP-seq. E) Clustering of all mC reader ChIP-seq peaks based on their colocalization. Five clusters were identified, which are bound by different subsets of mC readers, and show different chromatin environments as highlighted by DNA methylation levels in each context, the transcript abundance, and the presence of annotated gene and TE features. F) Browser displays of a representative locus for each cluster defined in (E). ChIP-seq enrichment (log_2_ Fold Change [IP/Input]) is shown for each mC reader as well as a WT control (where the IP was performed on a WT population, with no tagged protein), along with DNA methylation in each sequence context and RNA expression in WT seedlings (log_2_ CPM).

**Figure 3:**
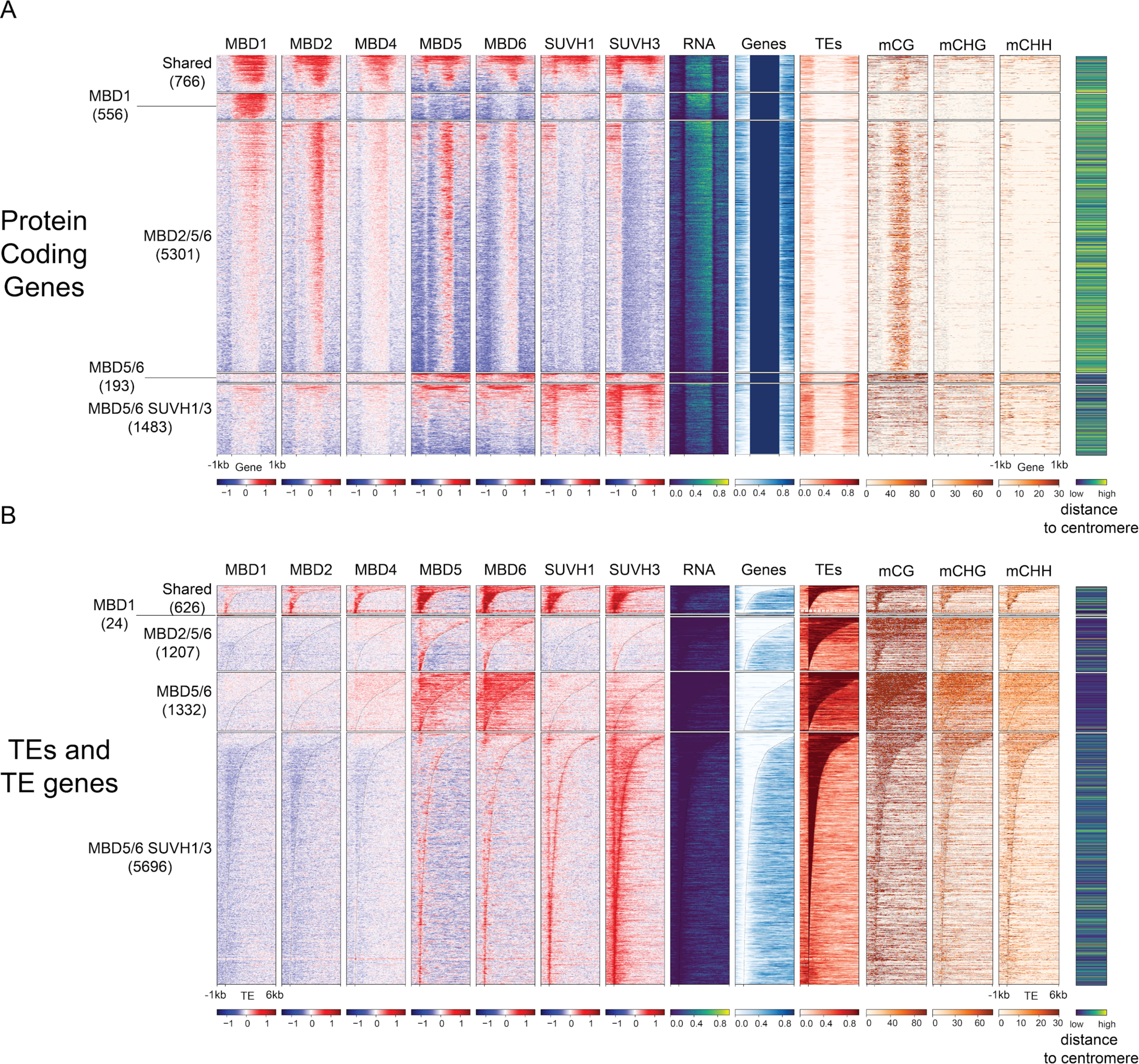
**mC readers differ in their binding preferences across genes and transposable elements.** A) Heatmaps of ChIP-seq signal enrichment of mC readers in protein-coding genes (TAIR10). Each cluster was sorted by descending means of the total read number in all bins of the region. For each locus, the methylation level in each context and the RNA expression level (normalized by RPKM) were calculated from Col-0 WT seedlings sample. For the methylation data, bins with no WGBS coverage are displayed in grey. ChIP-seq signal enrichments were calculated by comparing each ChIP sample to its corresponding input DNA control, scaled in log_2_FC from −1.5 to +1.5. The number of loci in each cluster are reported on the left hand-side of the heatmaps. The presence of an annotated gene or TE in each bin is shown in blue and red, respectively (0 if absent, 1 if present). Genes were scaled to 2kb, with 1kb upstream of the TSS and downstream of the TES, plotted in 100bp windows for DNA methylation tracks and 20bp windows for the others. B) Heatmaps of ChIP-seq signal enrichment of mC readers in transposable elements (TAIR10). Heatmap details as in panel A, except that TEs were aligned on their 5’ end, plotting 1 kb upstream and 6 kb downstream.

To more closely examine the similarities and differences of where these seven selected mC reader proteins were binding in the genome, we merged the peaks from each of the proteins into a non-overlapping set of regions, which were then clustered based on their ChIP-seq signal for each of the mC reader proteins (Fig 2E). Five major clusters were defined, with the first cluster corresponding to regions methylated in all sequence contexts, often intersecting annotated TEs, and bound by all seven mC reader proteins (Fig 2E,F). The second cluster comprises expressed, unmethylated genes, which are not bound by any mC reader, except for MBD1 (Fig 2E,F). The regions in the third cluster are bound by MBD2, MBD5, and MBD6, with these peaks corresponding to expressed genes that are methylated in the CG sequence context, reflective of GbM (Fig 2E,F). Looking specifically at all genes with GbM (Supp Fig 4) further confirmed this observation. The fourth cluster is only bound by MBD5 and MBD6, and is TE-rich and gene-depleted, highly methylated, and transcriptionally silenced (Fig 2E,F). Finally, the fifth cluster, bound by MBD5, MBD6, SUVH1, and SUVH3 is also TE-rich but surrounded by genes, potentially representative of shorter euchromatic TEs (Fig 2E,F). Taken together, this reveals that different mC readers have vastly different binding preferences, which are not only specific to the DNA methylation levels and sequence contexts, but to the more general chromatin state as well.

**Figure 4:**
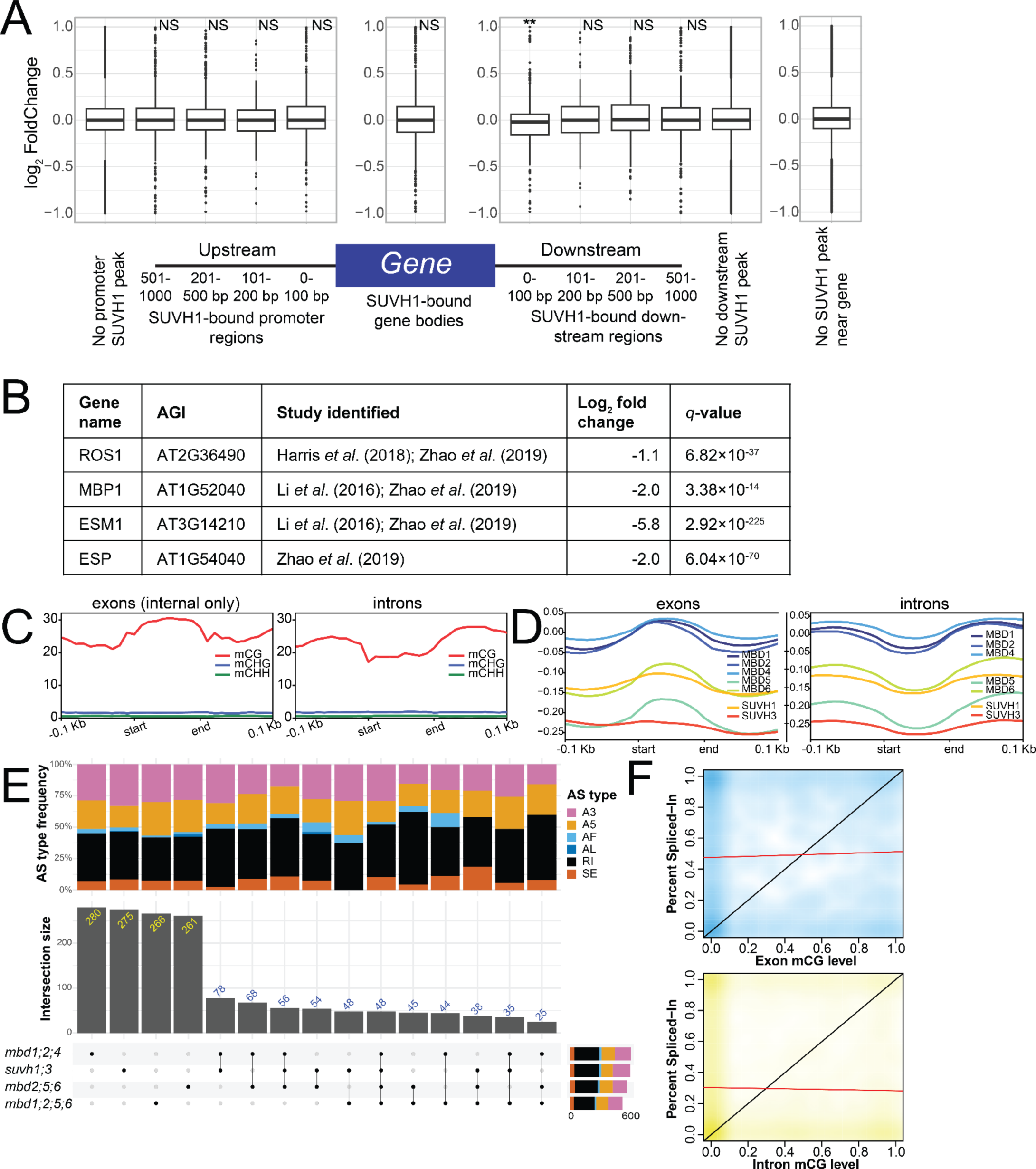
**Transcriptomic changes of higher order mC reader mutants.** A) Expression of protein-coding genes with and without SUVH1-binding sites at various locations relative to the transcriptional start site (TSS) and transcriptional end site (TES). SUVH1-bound gene bodies were compared to genes with no binding in the gene body or within 1 kb of the gene with regards to changes in expression in the *suvh1;3* mutant. The effect of binding within windows downstream of the gene on expression in the *suvh1;3* mutant were compared to genes without binding in the downstream region. The same was performed for those for binding upstream. NS = adjusted p-value > 0.05, ** = adjusted p-value < 0.01, after Benjamini-Hochberg correction of Mann-Whitney/Wilcoxon rank-sum test. B) Known targets of SUVH1/3 gene expression changes (log_2_FC and *q*-values) in our *suvh1;*3 double mutant. C) Metaplots of DNA methylation levels in each sequence context over exons and introns for the de novo transcriptome generated in this study. Exon/intron lengths were normalized to 100 bp using DeepTools. D) Metaplots of mC reader ChIP-seq (log_2_FC[IP/Input]) over exons and introns similarly to (C). E) Upset plot showing the number of differential alternative splicing events (DAS) shared between the different higher order mutants studied. F) Density plot of the correlation between exon (blue) or intron (yellow) mCG level and inclusion rate of the exon (or intron) within the final transcript, as measured by percent spliced-in (PSI). The black line in the below plots represents a linear relationship between methylation of the exon (or intron) and inclusion of the region in the final transcript and the red line shows the regression from a linear model of their relationship. The Spearman correlation coefficient for exon mCG and PSI is 0.02 (p-value = 0.4) and the Spearman correlation coefficient for intron mCG and PSI is −0.001 (p-value = 0.9).

To confirm that these patterns of mC reader binding throughout the genome are seen without the potential bias of peak calling being used to define the regions, we also focused on the ChIP-seq mapped read signal within protein-coding (genic) and TE regions (Fig 3). This showed that MBD1 is enriched in genic regions (Fig 3A), but is largely absent from TE sites (Fig 3B). Despite our initial probe-based approach identifying MBD1 as enriched in binding to methylated DNA (Fig 1), the ChIP-seq profiling of MBD1 reveals that, *in vivo*, MBD1 binds to regions both with and without mCG methylation, corresponding to unmethylated genes largely absent from the binding of other mC readers profiled here, with the exception of MBD2 at some methylated sites (Fig 3A). In contrast, MBD5 and MBD6 generally bind to the same regions as one another, which are defined by the presence of mCG, irrespective of whether it is a protein-coding gene or TE, and of whether it is expressed or not (Fig 3). MBD5 and MBD6 bound regions are often co-bound by MBD2, but only in protein-coding regions (Fig 3A), making genes with body methylation the only targets of MBD2 identified in this study. MBD4 binds to the fewest sites in the genome, to methylated sites bound by other mC readers (mainly MBD1/2/5/6) (Fig 3), as well as to unmethylated promoters (Fig 2A). SUVH1/3 were previously shown to bind to RdDM targets (Harris et al. 2018; Zhao et al. 2019). We confirmed this preference, and showed that SUVH1/3 targets are short euchromatic TEs and borders of larger heterochromatic ones (Fig 3B), located in the promoter of protein-coding genes (Fig 3A), with high levels of mCHH (Fig 2C,E).

Together, these analyses reveal that mC readers have specific binding specificities *in vivo*: MBD1 binds mostly poorly methylated genic regions; MBD2 binds almost exclusively to genes with body methylation; MBD5 and MBD6 bind to all mCG without discrimination, including in GbM-genes, short euchromatic TEs, and long heterochromatic TEs; and SUVH1 and SUVH3 bind to short euchromatic TEs (RdDM targets) that are highly enriched in non-CG methylation. Furthermore, these results suggest that the binding of different combinations of mC readers to a single genomic site may form its own code that might define or represent the epigenomic context of the region.

### Assessing the effects of the absence of multiple mC readers

Single and double mutants of some mC readers have previously been reported, as have some double and triple mutants, with only mild impacts on the transcriptome and epigenome (Harris et al. 2018; Ichino et al. 2021b; Zhou et al. 2021; Feng et al. 2021; Ichino et al. 2022). Given the co-binding of multiple mC readers to the same genomic sites within the Arabidopsis genome, we generated different combinations of mutants in two of the seven profiled mC readers by CRISPR/Cas9 editing (Supp Table 4). After confirming the presence of frame-shifting indels, these mutants were then crossed to produce higher order mutants, including the triple mutants *mbd2 mbd5 mbd6* (referred to as *mbd2;5;6*), *mbd1 mbd2 mbd4* (*mbd1;2;4*), and the quadruple mutant *mbd1 mbd2 mbd5 mbd6* (*mbd1;2;5;6*). We also generated a new line of the *suvh1 suvh3* (*suvh1;3*) double mutant. No morphological impact of these higher order mutants could be observed, so we explored their molecular phenotypes by performing RNA-seq and WGBS (Supp Fig 5). In totality, very few changes were found in either DNA methylation (Supp Fig 5A) or gene expression (Supp Fig 5B), the only exception being the *suvh1;3* double mutant, in which we detected over 2000 differentially expressed genes (Supp Fig 5B). Previously, SUVH1-binding in the promoter region immediately proximal to the TSS of genes was found to slightly enhance their expression (Harris et al. 2018). With the data generated here, we looked to see if loss of SUVH1 and SUVH3 (*suvh1;3*) (Supp Table 4) led to decreased expression of protein-coding genes with SUVH1-binding in their promoter regions, but detected no evidence of altered expression of these genes (Fig 4A, 3C). We confirmed that our *suvh1;3* double mutant was exhibiting the expected molecular phenotype by verifying that known targets of SUVH1/3 enhancement were down-regulated: *ROS1*, *ESM1*, *ESP* and *MBP1* (Fig 4B, 3D), (Li et al. 2016; Harris et al. 2018; Zhao et al. 2019). We did detect a small but significant reduction in the expression of genes where SUVH1 was bound immediately after the transcription end site (TES) (Fig 4A). These data suggest that this genome-wide effect is modest or experimental-condition specific, and that many of the differentially expressed genes in *suvh1;3* may be the results of changes in direct target dysregulation, such as of *ROS1*.

**Figure 5:**
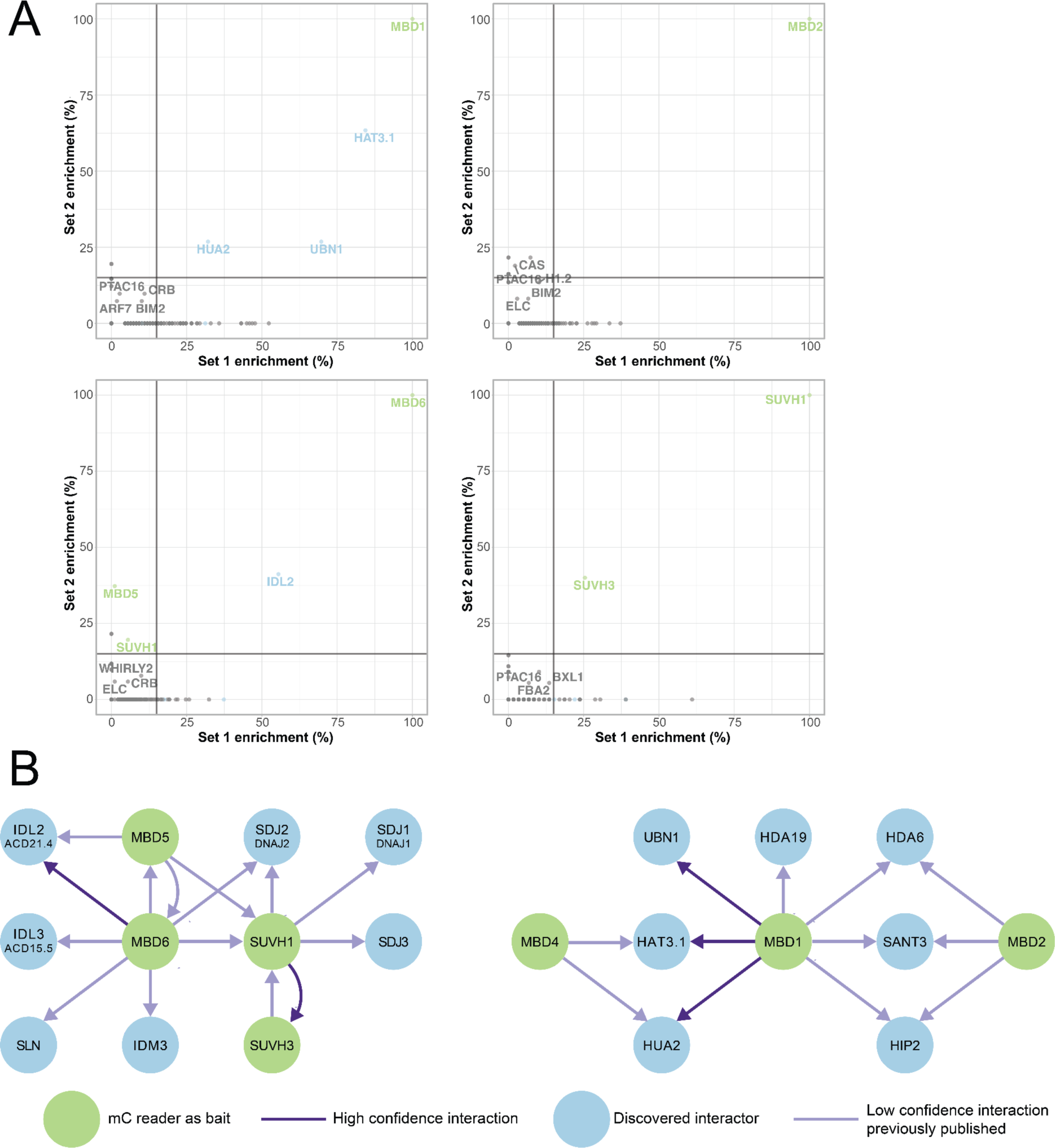
**Protein interactors with mC readers.** A) Scatter plots showing the correlation between Set 1 and Set 2 experiments for MBD1, MBD2, MBD6 and SUVH1. The grey lines represent the thresholds to reduce false positive proteins being selected for the interaction network. A 15% threshold for each replicate was selected. mC reader proteins that have been used as baits in this study are green. Interactors identified within this study are blue. B) A conservative interaction network of proteins in this study. Interactions (edges) identified in both Set 1 and Set 2 experiments are indicated in dark purple. Those edges that were present in our (T)AP but did not meet our strict criteria, but have been identified in another study are indicated in light purple. Nodes (proteins) are coloured as in panel A.

To investigate whether the CG-DMRs that we did identify (4446) were potentially linked to the loss of an mC reader, we overlapped the CG-DMRs with the mC reader ChIP-seq peaks, revealing little overlap between them (Supp Fig 5C), in agreement with previous work (Li et al. 2016; Harris et al. 2018; Zhao et al. 2019). To further investigate a potential role for an mC reader in directly controlling transcription and further investigate the potential role of GbM, we performed nuclear global run-on sequencing (GRO-seq) and 5’ GRO-seq (Hetzel et al. 2016) on the *mbd2* mutant, since MBD2 was specifically binding to GbM-genes. GRO-seq allows us to look beyond steady state RNA levels and to look at transcription initiation rates. We found few differences between WT and *mbd2* plants in terms of transcriptional activity as measured by GRO-seq and 5’ GRO-seq (Supp Fig 5D), and almost no overlap between the statistically different regions of transcription and MBD2 ChIP-seq peaks (Supp Fig 5E), suggesting that any differences were not resulting from the loss of a localised direct effect of MBD2 at these loci. Finally, we considered potential cross-talk between mC readers and histone modifications. Thus, we performed ChIP-seq in WT and mutant plants to examine a range of histone post-translational modifications associated with active (H3K27ac, H3K4me1, H3K4me2, H3K4me3, H3K36me3) and silenced (H3K9me2, and polycomb-deposited H3K27me3, H2AKub) chromatin states. Few differentially bound peaks were identified (Supp Fig 6A), and the binding profiles of most histone marks were highly similar between WT and each mutant examined (Supp Fig 6B). Taken together, these data suggest that loss of even multiple mC readers concurrently do not have profound effects on the DNA methylome, histone modifications, or transcriptome in Arabidopsis seedlings.

In many organisms, including flowering plants, mCG is enriched in exons relative to introns and has been linked to exon definition during RNA splicing (Zemach et al. 2010; Feng et al. 2010) (Fig 4C). We hypothesized that mC readers may act as adaptor proteins to recruit splicing factors to the chromatin (Luco et al. 2010; Pradeepa et al. 2012). We found that all mC readers examined except SUVH1 and SUVH3 were also enriched in exons compared to introns (Fig 4D), matching the enrichment of mCG levels, thus potentially supporting the notion that mC readers could recruit splicing factors to the chromatin during transcription. Further investigation, however, identified few (∼600) differentially alternatively spliced (DAS) events in the higher order mutants (Fig 4E), and there was very little overlap between higher order mutants with overlapping mutations (Fig 4E), suggesting that the few DAS events detected were likely background noise. Given the proposed role of DNA methylation within gene bodies in exon definition, we looked at inclusion levels of differentially spliced exons and introns in WT seedlings. If mCG was a mark for a region to be included in the final transcript, we would predict a correlation between exon (or intron) mCG level and inclusion rate of the exon (or intron) within the final transcript, as measured by percent spliced-in (PSI). Therefore, we correlated the WT mCG levels with the PSI values for all skipped exons and retained introns, but again found no correlation (Fig 4F). These data indicate that while mCG and many mC readers are enriched in exons, mCG does not appear to direct splicing inclusion of events in Arabidopsis seedlings, and mC readers do not direct alternative splicing.

### Identification of mC reader protein interactions

To gain insights into molecular mechanisms that the mC reader proteins may be involved in, protein interactors were identified by affinity purification coupled with mass spectrometry of the tagged mC readers used for ChIP-seq to identify the mC reader binding profiles. To ensure stringent identification of protein-protein interactions with mC reader proteins, one TAP-MS experiment was performed in one replicate (Set 1) and another AP-MS experiment was performed in triplicate (Set 2). TAP-MS for Set 1 was performed using the Strep and His tags on MBD1, MBD2, MBD6, and SUVH1 (Supp Table 3; Supp Table 5). Set 2 consisted of these same baits as well as MBD4, MBD5, and SUVH3, isolated using the Strep tag (Supp Table 3; Supp Table 5). To identify only the most robust interactions, we applied the following stringent rules: high-confident interactors that appear in the TAP-MS from both labs (Fig 5a), high confidence in the replicates from Set 2 experiment (Supp Table 5), or overlap with previously published interactions (Supp Table 6). Correlation of the peptide enrichment of putative interactors in each replicate revealed several protein interactors (Fig 5A). Notably, MBD1 pull-downs in both sets of experiments revealed interactions with histone modifying proteins UBN1, HAT3.1, and HUA2 (Fig 5). UBN1 is a component of the HIRA complex, responsible for histone variant H3.3 deposition (Nie et al. 2014). Other HIRA complex members (HIRA, UBN2, and H3.3) were also identified in Set 1 (Supp Table 6), and while not passing our stringent criteria, still suggest that MBD1 interacts with the HIRA complex. MBD1 also interacted with the transcription factors HAT3.1 and HUA2 (Fig 5b). HAT3.1 and HUA2 were also found to interact with MBD4 in Set 2. Another high-confidence interaction that we identified was between MBD6 and heat shock protein IDL2 (ACD21.4) (Fig 5). This interaction has been reported previously (Ichino et al. 2021a), as has the direct interaction of SUVH1 with SUVH3 that we identified (Fig 5) (Harris et al. 2018). A number of interactors that did not pass our initial stringent criteria (Supp Table 5) have been previously reported as mC reader interactors (Supp Table 6), thus they have been validated by another independent study and were included here in the network of mC reader interactors (Fig 5b). Such interactors include DNAJ proteins (SDJ2/DNAJ2, SDJ1/DNAJ1, and SDJ3) with SUVH1, and DNAJ proteins SLN and SDJ2/DNAJ2 with MBD6 (Fig 5b; Supp Table 6). We also found evidence for the interaction of MBD6 with heat-shock domain protein IDL3 (ACD15.5) and the de-methylation factor IDM3 (Fig 5b; Supp Table 6). Previous studies showed that MBD1, MBD2, and MBD4 interact with a histone deacetylase complex (Zhou et al. 2021; Feng et al. 2021). We found evidence for both MBD1 and MBD2 interacting with HDA6 and other proteins from this complex: SANT3 and HIP2 (Fig 5b; Supp Table 5; Supp Table 6), suggesting that MBD1 can take part in multiple complexes, one for regulating histone acetylation, and another for H3 variant deposition. Taken together, these data show that not only do the different mC readers have different binding profiles throughout the genome, but also form multiple different complexes, indicating a sub-functionalization of these factors.

## Discussion

Here, we used an unbiased screening approach to identify DNA methylation binding proteins in Arabidopsis and characterised the binding profiles of seven of these: MBD1, MBD2, MBD4, MBD5, MBD6, SUVH1, and SUVH3. While some of these mC readers had overlapping binding sites within the genome, clear differences between them could be identified. MBD2 was restricted to binding gene-body methylation, while MBD5/6 bound mCG sites within genes and TEs, often overlapping with SUVH1/3 binding sites, which largely bound non-CG methylation TE sites. Despite being identified within our screen as an mC reader, MBD1 is largely bound to unmethylated sites within gene bodies, which raises some interesting questions regarding what determines its *in vitro* versus *in vivo* binding profiles.

Despite the main focus of this study to be on mC readers, the methylated-probe pull-down screen performed has also identified proteins which preferentially bind unmethylated DNA (Fig 1). MBD9 has never been shown to bind DNA but instead histone H4 (Zemach and Grafi 2007; Yaish et al. 2009), and we found that MBD9 was strongly enriched with unmethylated DNA (Fig 1). The only protein enriched in two unmethylated contexts (CG and CHH) (Fig 1) was the AP endonuclease (AT2G41460), known to be involved with DNA repair post-active demethylation by DME or ROS1 (Córdoba-Cañero et al. 2011; Lee et al. 2014; Akishev et al. 2016), which along with the detection of the DNA repair protein RAD4 (AT5G16630) (Kunz et al. 2005; Liang et al. 2006), suggests that methylation sensitivity may play a role in this part of DNA repair. Two Jumonji-C (JmjC) domain-containing proteins, JMJ24 and JMJ28, were found enriched with unmethylated CHG (Fig 1). JMJ24 acts to reduce H3K9me2 levels indirectly, through ubiquitin E3 ligase activity via its RING domains that targets CMT3 for proteasomal degradation (Deng et al. 2015, 2016; Kabelitz et al. 2016). JMJ28 is an homolog of JMJ24 that also contains a JmjC and a RING domain (Lu et al. 2008). Our results support an association between JmjC-domain proteins and DNA methylation, especially in the CHG context, and might suggest a role in preventing heterochromatin marks from spreading into euchromatin like in other species, such as JmjC-domain proteins DMM-1 in *Neurospora crassa* and Epe1 in *Schizosaccharomyces pombe* (Tamaru 2010). Our comparison of *in vitro* (Fig 1) and our *in vivo* (Fig 2) binding preferences of the examined mC readers largely agreed with each other, with the exception of MBD1, which showed a preference for mCG *in vitro* but unmethylated DNA *in vivo*. This is likely explained by co-factors *in vivo* determining the binding profile of MBD1, but we do find MBD1 bound to some methylated sites *in vivo*, suggesting some variation in binding preferences within the genome. Like MBD5/6, MBD2 binds mCG in gene bodies, but unlike MBD5/6, MBD2 rarely binds TEs (Fig 2). MBD2 may interact with a range of chromatin remodelers, notably with BRAHMA (BRM) (Supp Table 5). BRM has been shown to prevent deposition of H3K27me3 (Li et al. 2015), which may aid in preventing repression of genes with GbM, helping to explain their constitutively expressed pattern (Coleman-Derr and Zilberman 2012). The recruitment of BRM to GbM would occur in addition to its recruitment by the H3K27 demethylase REF6. Given that REF6 preferentially binds unmethylated mCHG sites (Fig 1A,B), we propose a synergistic effect of REF6 and MBD2 might thus recruit BRM at genes with mCG but unmethylated CHG sites. Since GbM is absent from some plant species (Bewick et al. 2016), it would be interesting to know whether MBD2 presence and/or function is also conserved in these species, or if its involvement downstream of GbM has become moot.

Given the enrichment of many mC readers in exons relative to introns (Fig 4d), mirroring the reported patterns of CG methylation (Zemach et al. 2010; Feng et al. 2010) (Fig 4c), we explored whether alternative splicing could be directed by mC readers. Previously, histone modifications have been shown to direct alternative splicing in mammalian cells via adaptor proteins (Luco et al. 2010; Pradeepa et al. 2012). While an example of non-CG methylation in male sex cells of Arabidopsis is related to alternative splicing patterns (Walker et al. 2018; Lloyd and Lister 2022), our results suggest that GbM does not direct splicing patterns (Fig 4). This is supported by previous work in plants that compared DAS in WT to the *met1* derived epiRILs (Bewick et al. 2016). Other work has raised questions over whether exonic enrichment of DNA methylation is relevant to alternative splicing in insects (Patalano et al. 2015; Harris et al. 2019). Given the lack of an obvious effect on the loss of DNA methylation and mC readers on alternative splicing, it is likely that in many species, exonic DNA methylation is involved in a different mechanism, potentially as a by-product of CMT3 activity in flowering plants, as previously suggested (Bewick et al. 2016, 2017; Wendte et al. 2019), and that the MBDs examined here do not act in the “chromatin–adaptor complex” model of the regulation of alternative splicing (Luco et al. 2011).

The lack of obvious phenotypes within mC reader mutants, including higher order mutants, suggests that their role in normal plant growth is complex and masked by conventional techniques to study them. The lack of phenotype identified here with our higher order mutants is in contrast to Ichino *et al*. 2021, which found that the *mbd5;6* had defects in silencing (Ichino et al. 2021a). However, the use of floral tissue for their RNA-seq work would indicate a cell-type specific function for MBD5/6 (Ichino et al. 2022). The role in repression by MBD5/6 only appears to be detectable when normal chromatin compaction is lost, either naturally in pollen vegetative cells or via a mutation in the *H1* encoding genes (Ichino et al. 2021a, 2022). Considering the specialized function and regulation of DNA methylation in some tissues such as pollen vegetative cell and endosperm, it is likely that the loss of mC readers would only create phenotypes in these specific cell-types, or during restricted developmental stages. We found that loss of *suvh1;3* led to more transcriptomic changes than for any other higher order mutant studied here (Supp Fig 5) and found that previously reported targets of SUVH1/3-mediated expression enhancement, including *ROS1* (Harris et al. 2018; Zhao et al. 2019), were down-regulated in the *suvh1;3* double mutant (Fig 3D). Given that a similar number of genes were up- and down-regulated in *suvh1;3* plants, it is likely that many of these dysregulated genes are indirect targets of SUVH1/3 and might be the consequences of the loss of ROS1 and other targets. We showed here with both ChIP-seq and DAP-seq that MBD5 and MBD6 bind methylated CG in all genomic contexts (Fig 2,3; supp Fig 3). Also, no direct interactions with DDM1, MET1, or H1 were found in our TAP-MS (Fig 5, Supp Table 5) nor previously published datasets (Li et al. 2015, 2017; Ichino et al. 2021a). The mislocalization of MBD5/6 in *ddm1* mutants (Zemach et al. 2005) must then be due to the redistribution of mCG. MBD6 has also been shown to contribute to rDNA silencing in nucleolar dominance (Preuss et al. 2008). The role of these proteins might thus be to simply prevent transcription by steric hindrance, as the increase in accessibility in up-regulated genes in pollen grains might suggest (Ichino et al. 2022).

Our protein-protein interaction network for mC readers (Fig 5b) may indicate some mechanistic roles for these mC readers. The chromatin landscape in Arabidopsis is a complex interconnection of mechanisms that control gene expression, while maintaining TE silencing. We found that mC readers in Arabidopsis likely participate in this intricate regulation, binding to methylated sequences in specific chromatin contexts. It has been shown previously that MBD5 and MBD6 participate in gene silencing with SILENZIO (Ichino et al. 2021b, 2022). However, they also showed that MBD5 and MBD6 can interact with SUVH1 and SUVH3, proteins that participate in gene activation in Arabidopsis (Harris et al. 2018; Zhao et al. 2019). We have shown here that MBD5, MBD6, SUVH1, and SUVH3 are often colocalized in their genomic binding sites in seedlings, with the main difference being the preference for SUVH1/3 for non-CG methylation in contrast to CG methylation for MBDs. However, why some loci are bound by the reportedly activating SUVH1/3 mC readers (Harris et al. 2018; Zhao et al. 2019), and others by the repressive MBD5/6 mC readers (Ichino et al. 2021b, 2022), is unclear, especially given that the apparently repressive MBD5/6 proteins also bind to components of an activation complex containing IDM3 and SDJ2/DNAJ2 (Li et al. 2017) (Fig 5). It is conceivable that MBD5/6 might moonlight with both activator and repressor roles, depending on the exact chromatin context, such as whether they are in complex with SUVH1/3 or not (Fig 2 and 3). While IDM3 and SDJ1/2/3 have been reported to control DNA methylation levels (Miao et al. 2021), we did not see evidence for MBD5/6 or SUVH1/3 playing a role in this DNA methylation maintenance role (Supp Fig 5), which is consistent with previous reports of stable methylomes in mC reader mutants (Harris et al. 2018; Zhao et al. 2019; Ichino et al. 2021a). We found that MBD1 and MBD4 interacted with transcription factors HAT3.1 and HAU2, suggesting that these transcription factors may recruit MBD1/4 to specific sites in the genome, or that MBD1/4 may aid in providing specificity to HAT3.1/HAU2 with regards to the epigenomic context. The HIRA complex is responsible for the deposition of H3.3 histone variant (Nie et al. 2014). In addition to identifying HIRA complex component UBN1 as a high-confidence interactor of MBD1 (Fig 5), we also found HIRA, UBN2, and H3.3 as putative MBD1 interactors (Supp Table 5), suggesting that MBD1 may play a role in the recruitment of the HIRA complex. Previous reports have shown that MBD1/2/4 interact with a deacetylation complex composed of HDA6 and SANT-domain proteins and the TE-derived HHP1/HARB (Zhou et al. 2021; Feng et al. 2021). Feng *et al*. (2021) found that loss of MBD1, MBD2, and MBD4 lead to a small change in H3 acetylation by ChIP-seq (Feng et al. 2021), and while we did observe a slight decrease in H3K27ac in *mbd1* and *mbd1;4*, we did not examine the triple *mbd1;2;4* mutant as they did (Supp Fig 6). Also, we used an antibody to a specific acetylation mark (H3K27ac) rather than a general antibody to any H3 acetylation, as performed in (Feng et al. 2021). The exact factors and features that determine the *in vivo* preference of MBD1 for unmethylated DNA (Fig 2) but the *in vitro* preference for methylated DNA (Fig 1) are unclear but could be related to the multiple proteins that our data suggest can interact with MBD1: the HIRA complex (Fig 5), the histone deacetylation complex (Zhou et al. 2021; Feng et al. 2021) (Fig 5), and binding with transcription factors HAT3.1 and HUA2 (Fig 5). MBD2 almost exclusively binds CG methylated gene bodies (Fig 2). The co-binding of MBD1 and MBD2 to the HDA6 complex (Zhou et al. 2021; Feng et al. 2021) (Fig 5) may indicate that this complex interacts with the fraction of MBD1 bound to methylated regions of the genome, whereas MBD1-specific complexes, notably the HIRA complex (Fig 5), may be recruited to unmethylated regions of the genome. It has been recently shown that the chromatin remodeler DDM1 is only able to facilitate histone H3.1 deposition in nucleosomes containing unacetylated histone H4, thus allowing for DNA methylation to take place (Lee et al. 2023). MBD1, MBD2 and MBD4 might then participate in this interplay between DNA methylation, histone acetylation, and histone variant deposition, crucial for epigenetic inheritance (Lee et al. 2023). The lack of a clear phenotype, even in the higher order mutants, raises questions regarding the function of mC readers in plants. The GbM binding profile, exemplified by MBD2, could indicate a protective role for this protein against DNA damage. DNA methylation is mutagenic, but genes with GbM have a lower rate of evolution (Takuno and Gaut 2012), suggesting that a mechanism exists that prevents mutations of these cytosines. Confirming this hypothesis would require a much longer time-frame and many generations to see consequences of mutations.

Taken together, we have shown that mC readers form multiple protein complexes, which suggest sub-functions within the different mC reader groups that relate to their specific binding profiles within the Arabidopsis genome. These results support the existence of independent regulatory mechanisms downstream of DNA methylation that function in a chromatin context dependent manner. Further investigation, in specific cell-types or in plants deficient in epistatic mechanisms, could be key to address the apparent redundancy and to understand the precise function of these proteins.

## Material and methods

### DNA affinity pull-down assay

#### Nuclear protein extraction from Arabidopsis cell culture

Arabidopsis Landsberg erecta (Ler) ecotype cell suspensions derived from root callus was cultured in growth media (1x Linsmaier and Skoog basal salts with minimum organics, 3% (w/v) sucrose, 0.5 mg/L naphthalene acetic acid and 0.05 mg/L kinetin, adjusted to pH 5.7 with KOH) at 25°C under constant darkness and 130 rpm orbital shaking. Cultures were maintained by inoculating 20 mL of 7-day-old cells into 100 mL of fresh media in 250 mL Erlenmeyer flasks. Cultured cells were pelleted for 10 min at 500 rpm and protoplasted by enzymatic digestion in Enzyme Buffer (0.4 M mannitol, 3% sucrose, 8 mM CaCl2, 1% cellulase, and 0.5% macerozyme) for 4 hours. The protoplast solution was then filtered through 70 µM nylon mesh and centrifuged for 10 min at 26 xg after addition of 1 volume Mannitol/W5 Buffer (0.4 M mannitol and 0.2x W5 where 1x W5 is 5 mM glucose, 154 mM NaCl, 125 mM CaCl_2_, 5 mM KCl, and 1.5 mM MES, pH adjusted to 5.6 with 0.1 M KOH). Pelleted protoplasts were washed twice with Mannitol/Mg Buffer (0.4 M mannitol, 0.1% MES and 15 mM MgCl_2_, pH adjusted to 5.6 with 0.1 M KOH), resuspending the protoplasts by gentle rocking and centrifuging for 10 min at 26 x g. Pelleted protoplasts were resuspended in Nuclei Isolation Buffer (NIB: 20% glycerol, 10 mM Tris-HCl pH7.5, 10 mM MgCl_2_, 10 mM KCl, 0.5% triton X-100, 10 mM β-mercaptoethanol and 1x EDTA-free complete protease inhibitors), filtered through two sheets of miracloth and pelleted at 1500 x g for 20 min. Pellets were resuspended in NIB and centrifuged at 800 x g for 15 min. Nuclei were resuspended in Nuclei Resuspension Buffer (50% glycerol, 50 mM HEPES-KOH pH 7.6, 100 mM NaCl, 5 mM MgCl_2_, 10 mM KCl, 1 mM DTT, and 1x EDTA-free complete protease inhibitors) for visualization under fluorescence microscope by DAPI staining, or nuclear proteins were extracted by resuspending the nuclei in Nuclei Extraction Buffer (20 mM Tris-HCl (pH 7.9), 420 mM KCl and 1.5 mM MgCl_2_, 0.5 mM DTT, and 1x EDTA-free complete protease inhibitors), filtering the extract through a 12-gauge needle followed by centrifugation to pellet the membrane and debris. Protein concentration in the supernatant was measured with the Bio-Rad Protein Assay, establishing a linear regression with a range of bovine serum albumin standards.

#### Probes design

DNA probes were designed for each sequence context following the model of (Spruijt et al. 2013). For each possible 5 bp motif containing a unique cytosine in each context (i.e. in the CG context, all WCGWW motifs, where W is A or T), the number of sites in the Arabidopsis genome were evaluated, grouped by genomic feature, and the average methylation of the cytosine was calculated from Col-0 WT bisulfite sequencing data (Secco et al. 2015). The motifs found with higher frequency in more highly methylated annotated sequences (genes and TEs) were selected. The motifs were repeated 4 times on the probe and flanked by unmethylated sequences to create the 33-bp oligos (Table 1).

**Table 1.**
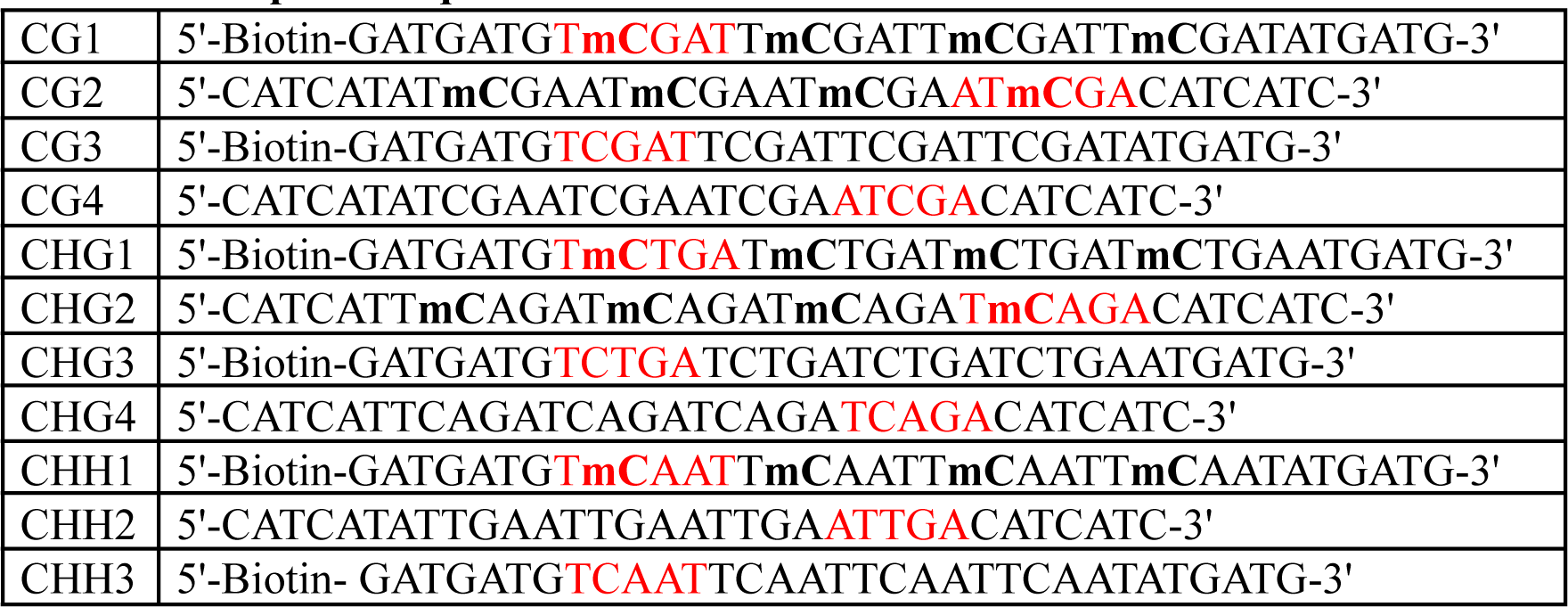
DNA probe sequences and modifications.

DNA probes (Table 1) were synthesized as single-stranded oligonucleotides by IDT, to be paired as follows: CG1 and CG2 constitute the methylated CG probe, and CG3 and CG4 the unmethylated CG control probe; CHG1 and CHG2 constitute the methylated CHG probe, and CHG3 and CHG4 the unmethylated CHG control probe; CHH1 and CHH2 constitute the CHH methylated probe, and CHH3 and CHH2 the unmethylated control probe. DNA affinity pull-down assays were then performed following an adapted version of (Spruijt et al. 2013). Oligos were paired by incubating 20 µg of each oligo in NEB buffer at 95°C for 5 min and gradually cooled-down overnight. Then, 10 µL of streptadivin sepharose high-performance beads (GE Healthcare) were used to immobilize the DNA probes, verifying that the probes were indeed immobilized on the beads by agarose gel electrophoresis. Three technical replicates were performed for each probe. For each replicate, 450 µg of nuclear protein extract was incubated with the DNA probes, in presence of 10 µg of dAdT as competitor DNA, rotating for 90 min. Beads were washed twice with PBS buffer and proteins bound to the probes were digested with trypsin. Tryptic peptides were eluted in Peptide Elution Buffer (100 mM Tris-HCl pH 7.5, 2 M urea, and 10 mM DTT) and loaded on C18 Stage-Tips pre-activated by methanol, washed once with Buffer B (0.1% formic acid, 80% acetonitrile) and washed twice with Buffer A (0.1% formic acid). Samples were then washed once with Buffer A.

#### LC-MS/MS measurements and statistical analysis

Peptides were eluted from Stage-Tips in Buffer B, after which the acetonitrile was evaporated using a vacuum concentrator. The samples were resuspended in 12 μl of buffer A, of which 5 μl were loaded onto a 30 cm column packed with 1.8 μm Reprosil-Pur C18-AQ (Dr Maisch GmbH). A 114 min gradient of acetonitrile (7%-32%), followed by washes at 50% then 90% acetonitrile, was employed to elute the peptides from the column, using an Easy-nLC 1000 system (Thermo Fisher Scientific). Eluted peptides were sprayed directly into an LTQ-Orbitrap Fusion Tribrid mass spectrometer (Thermo Fisher Scientific). Scans were collected in data-dependent top-speed mode of a 3-second cycle with dynamic exclusion set at 60 sec, for 140 min of total data collection.

Measured peptides were searched against the UniProt Arabidopsis thaliana proteome (version 2014-09-03) with MaxQuant version 1.5.1.0 (Cox and Mann 2008). Default settings were used, with additional options for ‘match between runs’, ‘label free quantification (LFQ)’ and ‘intensity based absolute quantification (iBAQ)’ enabled. Data were analysed with Perseus version 1.4.0.0 and in-house R scripts. Reverse and contaminant hits were removed, and the LFQ columns were transformed into log_2_ scale. Resulting data were filtered for proteins with three valid values in at least one of the samples, after which missing values were imputed by semi-random, low values (width = 0.3, shift = 1.8). To generate volcano plots between two samples, statistical outliers were determined using a two-tailed *t*-test with a permutation-based false discovery rate (FDR). The following different FDR and s0 (similar to a minimal fold-change) cut-offs were used to limit the number of proteins enriched in each context: CG: FDR=0.005, s0=3.0; CHG: FDR=0.025, s0=5.0; CHH: FDR=0.05, s0=4.0. To generate a hierarchical clustering, an ANOVA was executed with all the samples, also with a permutation-based FDR. Insignificant proteins were discarded, then the median of each row was subtracted to get deviations from the median. Statistical analysis and original plots were generated with Perseus software (Tyanova et al. 2016). R was used to generate the final plots.

### DNA Affinity Purification Sequencing (DAP-seq)

DAP-seq has been performed according to the original protocol (Bartlett et al. 2017). Genomic DNA libraries were created using genomic DNA extracted from 2-week old Col-0 seedlings grown on ½ Murashige and Skoog media supplemented with 1% sucrose with DNeasy Plant Mini kit (Qiagen). Plasmids containing the proteins of interest in the pIX-HALO backbone were ordered from ABRC, and an empty pIX-HALO plasmid was used as negative control (Table 2).

**Table 2.** Plasmids containing the proteins of interest in the pIX-HALO backbone.

|  |  |
| --- | --- |
| MBD1 | HALO TF-13 H08 |
| MBD2 | HALO TF-18 D10 |
| MBD4 | HALO TF-18 H12 |
| MBD5 | HALO TF-13 B06 |
| MBD6 | HALO TF-19 D09 |
| SUVH1 | HALO TF-20 B11 |
| SUVH3 | HALO TF-19 D07 |
| Empty | CD3-1742 |

Proteins were expressed using the TnT Coupled Wheat Germ Extract System with SP6 promoter (Promega). Protein expression was confirmed by Western Blot. After incubation of the proteins with DAP and ampDAP libraries, DAP-seq and ampDAP-seq libraries were pooled, separated by agarose gel electrophoresis, and the 200 to 400 bp range was purified by gel extraction using Isolate II PCR and Gel Kit (Bioline). Libraries were sequenced (100 bp paired-end) on a HiSeq 1500 instrument (Illumina) according to the manufacturer’s instructions. Raw sequencing data were then de-multiplexed with bcl2fastq software (Illumina).

### Plant material

#### Collection of single and higher order mutants

All plants were Arabidopsis thaliana accession Col-0, grown on soil at 22°C in 16h light/8h dark cycles. For SUVH1, SUVH3, MBD1, MBD2, and MBD6, existing T-DNA lines were ordered from the Arabidopsis Biological Resource Center (Supp Table 4). For MBD4 and MBD5, single mutants were created by CRISPR/Cas9 mutagenesis. Guide RNAs were designed for each target using CRISPRdirect (Naito et al. 2015) and further analyzed with E-CRISPR (Heigwer et al. 2014). The cassettes containing two gRNAs for each mutant pair were ordered as gBlocks (IDT) and were inserted into pHEE2E backbone, provided by Dr Qi-Jun Chen (Wang et al. 2015), by restriction-ligation using BsaI (NEB). WT plants were transformed by Agrobacterium-mediated T-DNA insertion using the floral dipping procedure (Clough and Bent 1998). Plants transformed with the CRISPR cassette were selected by resistance to hygromycin. Genomic DNA was extracted from leaf tissue (Edwards et al. 1991). For T-DNA lines, genotyping was performed via PCR amplification of the inserted T-DNA sequence. The primers used for genotyping PCRs were designed with the iSect tool from the Salk Institute Genomic Analysis Laboratory and are listed in (Supp Table 7). Plants homozygous for the insertion were selected for subsequent experiments.

For CRISPR/Cas9 mutated plants, homozygous plants were determined by the presence of a single trace including a mutation after Sanger sequencing (primers used are listed in Supp Table 7). Only homozygous T2 plants that were selected for the absence of the CRISPR cassette by PCR were selected for subsequent experiments. Higher order mutants were generated by crossing the wanted mutations (Supp Table 4).

#### Complementation of mutants with epitope-tagged proteins under endogenous promoter

For each candidate, the single mutant line from T-DNA or CRISPR/Cas9 described above was complemented with an epitope-tagged version of the respective protein driven by their endogenous promoter. Promoters were defined by incorporating the closest DNase I hypersensitivity (DHS) sites upstream of the transcription start site, from DHS data generated from 7-day old seedlings in Col-0 background (Sullivan et al. 2014). 3’-UTRs were included for MBD6, as a DHS site was also present downstream, near the transcription termination site. Promoters and 3’ UTRs were amplified from Col-0 genomic DNA using the primers listed in (Supp Table 3) with Q5 Hot Start High-fidelity DNA polymerase (NEB). For MBD5 and SUVH3, Gibson Assembly was performed to insert these fragments into the backbone. For the others, PmeI (or SacI) and AvrII restriction sites were incorporated at the 5’ end of the primers to allow for restriction-ligation of the promoters into plasmids containing the SHH tag (2xStrep-HA-6xHis), generously provided by Dr. Dmitri Nusinow. Insert and entry plasmids were digested with PmeI (or SacI) and AvrII enzymes (NEB). Generated fragments were gel extracted, ligated and purified ligation products were electroporated in TOP10 *Escherichia coli* competent cells. Transformed bacteria were selected by resistance to spectinomycin and plasmid sequences were confirmed by Sanger sequencing. For MBD6, 3’-UTRs were then incorporated through a similar process using EcoRV restriction sites. Final plasmids were inserted into GV3101 *Agrobacterium tumefaciens* by heat-shock for plant transformation. T1 plants were grown on soil and transformants selected by resistance to glufosinate. Presence of both the original mutation and the complementation plasmid was confirmed by PCR using primers listed in Supp Table 7. To assess the expression level of the complemented plasmids, RNA was extracted from a population of T2 seedlings grown on ½ Murashige and Skoog media supplemented with 1% sucrose for two weeks with TRIzol Reagent (ThermoFisher Scientific). RNA was treated with RQ1 DNase (Promega), purified and converted to cDNA with sensiFAST cDNA Synthesis Kit (Bioline), using the provided mix of random hexamers and anchored oligo dT primers. qPCR reactions were performed with KAPA SYBR FAST qPCR Kit (Sigma-Aldrich) on a LightCycler 480 instrument (Roche). The qPCR primers used for this study are listed in Supp Table 7.

### Chromatin immunoprecipitation sequencing (ChIP-seq)

Three to eight grams of seedlings grown on ½ Murashige and Skoog media supplemented with 1% sucrose were harvested 14-days after germination induction and crosslinked in Crosslinking Buffer (10mM HEPES-NaOH, pH 7.4; 1% Formaldehyde) by drawing vacuum for 10 min, twice. Seedlings were transferred to Quenching Buffer (10mM HEPES-NAOH, pH 7.4; 200 nM Glycine) and vacuum applied for 10 min. After 3 washes in ddH_2_O, seedlings were frozen in liquid N_2_ and stored at −80°C. Tissue was grounded to fine powder using a mortar and pestle in liquid N_2_ and resuspended in Buffer A (10mM Tris-HCl, pH 8.0; 400mM Sucrose; 5mM β-Mercaptoethanol; 1x Protease inhibitors), rotating for 10 min. Samples were kept at 4°C from this step up to the reverse crosslinking step, including during the centrifugation steps. After filtering through a sheet of miracloth, samples were centrifuged for 20 min at 2,880 x g. Pellets were resuspended in Buffer B (10 mM Tris-HCl, pH 8.0; 250 mM Sucrose; 10 mM MgCl_2_; 1% Tx-100; 5 mM β-Mercaptoethanol; 1x Protease inhibitors) and centrifuged for 10 min at 12,000 x g. Pellets were then resuspended in Buffer C (10 mM Tris-HCl, pH8.0; 1.7 M Sucrose; 2 mM MgCl2; 0.15% Tx-100; 5 mM B-Mercaptoethanol; 1x Protease inhibitors) and overlaid on Buffer C cushion before centrifuging for 1h at 16,000 x g. Pellets were washed twice in Wash Buffer (10 mM Tris-HCl, pH 8.0; 200 mM NaCl; 1 mM EDTA, pH 8.0; 0.5 mM EGTA, pH 8.0; 1x Protease inhibitors) by resuspending in the buffer and centrifuging for 5 min at 4,000 x g. Final pellets were resuspended in Shearing Buffer (10 mM Tris-HCl, pH 8.0; 1 mM EDTA, pH 8.0; 0.1% SDS; 1x Protease inhibitors) and sonicated with Covaris S2 (12 min, duty cycle 5%, intensity 4, 200 cycles per burst, peak power 140). Triton X-100 and NaCl were added to the samples to reach a final concentration of 1% and 150mM, respectively. Chromatin was then cleared of debris by centrifuging for 10 min at 10,000 x g.

Chromatin (50 µL; 5%) was put aside to be used as input control. Then 4 µg of primary antibodies (αHA antibody: Biolegend, #901502; αH2A.Z antibody: abcam, ab4174; αH2AK121ub antibody: Cell Signaling Technology, 8240S; αH3K4me1 antibody: abcam, ab8895; αH3K4me2 antibody: abcam, ab32356; αH3K4me3 antibody: abcam, ab8580; αH3K36me3 antibody: abcam, ab9050; αH3K27me3 antibody: abcam, ab6002; αH3K27ac antibody: abcam, ab4729; αH3K9me2 antibody: abcam, ab1220; αH3 antibody: abcam, ab1791) were added to 1 mL of chromatin and incubated slowly rotating overnight. For each sample, 25 µL Protein G beads (Life Technologies) and 25 µL Dynabeads M-280 sheep anti-mouse IgG (Thermo Fischer Scientific) were equilibrated by washing three times with IP Buffer (10 mM Tris-HCl, pH 8.0; 1 mM EDTA, pH 8.0; 0.1% SDS; 1% Triton X-100; 150 mM NaCl) prior to add the chromatin, and samples were incubated rotating for 90 min. Protein/DNA complexes bound to the beads were washed twice with Low Salt Wash Buffer (20 mM Hepes-KOH, pH 7.9; 2 mM EDTA; 0.1% SDS; 1% Triton X-100; 150 mM NaCl), twice with High Salt Wash Buffer (20 mM Hepes-KOH, pH 7.9; 2 mM EDTA; 0.1% SDS; 1% Triton X-100; 500 mM NaCl), once with LiCl Wash Buffer (100 mM Tris-HCl, pH 7.5; 0.5 M LiCl; 1% NP-40; 1% Sodium Deoxycholate) and once with TE/10 Buffer (10 mM Tris-HCl, pH 8.0; 0.1 mM EDTA). Immunoprecipitated samples were resuspended in 49 µL Proteinase K Digestion Buffer (20 mM HEPES, pH 7.9; 1 mM EDTA; 0.5% SDS) and 1 µL of 20 mg/mL Proteinase K was added to each sample. In parallel, 2 µL of 10% SDS and 1 µL of 20 mg/mL Proteinase K was added to the input samples. Both types of samples were then incubated at 50°C for 15 min. DNA/protein complexes eluted from the beads were then transferred to a new tube. Subsequently 3 µL 5M NaCl and 0.5 µL 100 mg/mL RNase A were added to immunoprecipitated and input samples and were incubated overnight at 65°C shaking at 1,000 rpm to reverse the cross-linking. The next day, an additional 1.5 µL 20 mg/mL Proteinase K were added to all samples and samples were incubated for 60 min at 50°C. DNA was finally purified with AMPure beads and 15 µL of each sample were recovered. Input DNA (2 ng) and immunoprecipitated samples (5 µL) were processed according to Rubicon thruPLEX DNA-seq kit, using half reactions per sample. Libraries were sequenced on a HiSeq 1500 instrument (Illumina) according to the manufacturer’s instructions. Raw sequencing data were then de-multiplexed with bcl2fastq software (Illumina).

### Affinity purification coupled to mass spectrometry

Three to eight grams of seedlings grown on ½ Murashige and Skoog media supplemented with 1% sucrose were harvested 14-days after germination induction, frozen in liquid N_2_ and stored at −80°C. Tissue was grounded to fine powder using a mortar and pestle in liquid N_2_ and resuspended in Extraction Buffer (2 M hexylene glycol; 20 mM PIPES-KOH, pH 7.0; 10 mM MgCl_2_; 5 mM β-mercaptoethanol), rotating for 10 min. Samples were kept at 4°C from this step until the trypsin digestion, including during the centrifugation steps. After filtering through a sheet of miracloth, triton X-100 was added in small aliquots until the final concentration reached 1%. Samples were overlaid on a density gradient of 30% and 80% Percoll Buffer (2 M hexylene glycol; 5 mM PIPES-KOH, pH 7.0; 10 mM MgCl_2_; 1% triton X-100; 5 mM β-mercaptoethanol; 30 or 80% percoll) and centrifuged for 30 min at 2,000 x g. Nuclei were collected from the 30/80% percoll interphase, underlaid with 30% Percoll Buffer and centrifuged for 10 min at 2,000 x g. Pelleted nuclei were resuspended in SII Buffer (100 mM Sodium Phosphate, pH 8.0; 150 mM NaCl; 5 mM EDTA; 5 mM EGTA; 0.1% triton X-100; 1x protease inhibitors; 1x phosphatase inhibitors; 1 mM PMSF; 50 µM MG-132), and membranes were disrupted by sonication (40% power, 1s on/1s off, for 20s in total, repeated 3 times per sample). Samples were clarified by two successive centrifugation steps at 14,000 x g for 10 min.

For the experimental Set1, IBA MagStrep “type3” XT beads (Fisher Biotech) were washed twice in SII Buffer before adding the protein extract and were then incubated rotating for 60 min. The beads were washed twice in SII Buffer and three times with Strep-to-His Buffer (100 mM Na_2_HPO_4_/NaH_2_PO_4_, pH 8.0; 150 mM NaCl; 0.05% triton X-100). Protein complexes were eluted twice with Strep Elution Buffer (100 mM Na_2_HPO_4_/NaH_2_PO_4_, pH 8.0; 150 mM NaCl; 0.05% triton X-100; 50 mM biotin) by incubating rotating for 10 min. Dynabeads His-Tag Isolation and Pulldown (Thermo Fisher Scientific) were washed twice in Strep-to-His Buffer before adding the eluted proteins and incubating while rotating for 20 min. The beads were washed twice with Strep-to-His Buffer and three times with 25 mM ammonium bicarbonate before being snap-frozen in liquid N_2_ and stored at −80°C. Beads were resuspended in 50 µL Protein Elution Buffer (100 mM Tris-HCl, pH 7.5; 1 M Urea; 10 mM DTT) and incubated for 20 min, shaking. 5 µL 0.55 M iodoacetamide were added to each sample and the tubes were incubated for 10 min, shaking in the dark. 2.5 µL of trypsin (0.4 µg/µL in 0.01% TFA) were added to each sample and the tubes were incubated for 2h, shaking. After transferring the eluted peptides to a new tube by immobilizing the beads with a magnet, 50 µL were added to the beads for a second elution, after incubating for 5 min shaking. Then 1 µL of trypsin was added to the combined eluates and the tubes were incubated while shaking overnight. Peptides were loaded on C18 resin MicroSpin Columns Silica C18 (The Nest Group) which were pre-activated by methanol, washed once with Buffer B (0.1% formic acid, 80% acetonitrile) and washed twice with Buffer A (0.1% formic acid). Samples were then washed once with Buffer A and eluted from the Stage-Tips in Buffer B, after which the acetonitrile was evaporated using a vacuum concentrator. Samples were resuspended in 30 μl of 5% (v/v) acetonitrile/0.1% (v/v) formic acid before online reversed phase nanoflow (EASY-Spray HPLC column 50 cm x 75 µm ID) ESI coupled to an Orbitrap Fusion (ThermoFisher). Gradients were 5–35% (v/v) acetonitrile in 0.1% (v/v) formic acid (250 nl min−1) formed by a Dionex UltiMate 3000 series HPLC (ThermoFisher) over 120 min. Spectra were acquired in data dependent mode with a 120k resolution survey scan from 300–1500 m/z followed by selection of the eight most abundant doubly or triply charged ions for MS/MS analysis. Ions were dynamically excluded for 60 sec. Raw data was converted to mzML format by msconvert (3.0.9992) before spectral matching against the Araport11 peptide sequence database by CometMS (2016.01 rev. 2). Results were processed through the Trans-Proteomic Pipeline (5.0) tools peptide and protein prophet. Proteins with a probability above 0.9 were accepted for further analysis. For the experimental Set2, single strep IP protein pull-downs using 200 μg input lysate were also performed. We note that the MS/MS ID rates for these protein pull-downs are very low, for reasons we have not been able to decipher.

### Whole genome bisulfite sequencing (MethylC-seq)

Genomic DNA was extracted from 2-week old seedlings with DNeasy Plant Mini Kit (Qiagen) and further purified with Isolate II Gel and PCR Clean-up Kit (Bioline). Libraries were then generated from fragmented DNA (Covaris, 250 bp) with NxSeq AmpFREE Low DNA Library Kits (Lucigen). After size selection by AMPureXP beads, bisulfite conversion was performed with EZ DNA Methylation Gold kit (Zymo Research). Libraries were amplified with Kapa HiFi HotStart Uracil+ ReadyMix (Roche) and purified with AMPureXP beads. MethylC-seq libraries were sequenced on HiSeq 1500 platform (Illumina) using single-end 100 bp format, according to the manufacturer’s instructions.

### High-throughput RNA sequencing (RNA-seq)

Three biological replicates were performed in parallel for all genotypes from populations of 2-week old seedlings grown on ½ Murashige and Skoog media supplemented with 1% sucrose. Total RNA was extracted with RNeasy Plant Mini Kit (Qiagen) and treated with RQ1 DNase (Promega). Libraries were then generated with TruSeq Stranded Total RNA Library Prep Kit (Illumina), after depletion of ribosomal RNAs with Ribo-Zero rRNA Removal Kit Plant (Illumina). RNA-seq libraries were sequenced on a HiSeq 1500 (Illumina) using paired-end 42 bp format, according to the manufacturer’s instructions.

### Global run-on sequencing (GRO-seq)

Global Run-On sequencing (GRO-seq) and 5’-GRO-seq were performed on 10-20g of 2-week old seedlings (∼10 plates), according to the published protocol (Hetzel et al. 2016).

### Data analysis

#### mC reader ChIP-seq and (amp)DAP-seq

Illumina adapters were trimmed from the raw data using cutadapt (Martin 2011) with default parameters. Remaining reads were mapped to the Arabidopsis TAIR10 genome reference using bowtie2 in default end-to-end mode (Langmead and Salzberg 2012). Resulting files were converted to sorted and indexed bam files using the samtools suite (Li et al. 2009). Peaks were called with the version 2 of MACS (Zhang et al. 2008) by comparing each immunoprecipitated sample to its input, including for a WT sample (Col-0 plant with no tagged protein). For DAP and ampDAP-seq, the samples were compared to the DNA control library before and after amplification, respectively.

For each ChIP sample, peaks with 10% reciprocal overlap with the peaks called in WT were discarded, and to increase stringency, only the remaining top 80% based on local fold-change constitute the final set of peaks for each sample. These sets were merged into a single file by merging 50-bp overlapping peaks to generate a list of unique bound loci, using `bedmap –echo-map-range –echo-map-id-uniq` (Neph et al. 2012) and bedtools merge (Quinlan and Hall). These loci were annotated with annotatePeaks.pl from the Homer suite (Heinz et al. 2010), and then intersected with TAIR10 annotated transposable elements and transposable element genes with bedtools intersect. The deepTools suite (Ramírez et al. 2016) was used to convert bam files to bigwig files and to plot the heatmaps and profiles. The other figures were generated using R code based on the ggplot2 package (Wickham 2016) and UpSetR package (Conway et al. 2017). The homer suite was used to identify motifs under the ChIP-seq peaks (Heinz et al. 2010).

#### Histone modification ChIP-seq

Illumina adapters were trimmed from the raw data using cutadapt (Martin 2011) with default parameters. Remaining reads were mapped to Arabidopsis TAIR10 genome using bowtie2 in default end-to-end mode (Langmead and Salzberg 2012). Resulting files were converted to sorted and indexed bam files using the samtools suite (Li et al. 2009). Peaks were called with the version 2 of MACS (Zhang et al. 2008) by comparing each immunoprecipitated sample to its input. Peaks called for each histone modification from each genotype (WT and mC reader mutants) were merged with the following command: `cat WT_Mark.narrowPeak MutantGenotype_Mark.narrowPeak | sort −k1,1 −k2,2n | bedtools merge`, with mark being the histone modification in question, and MutantGenotype being whatever mutants were used to study alterations in that histone modification. Then edgeR (Robinson et al. 2010; Nikolayeva and Robinson 2014) was used to test for differential binding between WT and each mC mutant for each histone modification, using the merged histone peak files as a reference.

#### MethylC-seq

Raw sequencing data were base-called and de-multiplexed with the bcl2fastq software (Illumina). FASTQ files were mapped to the TAIR10 genome (previously processed to a 3-letter genome reference) with BS-Seeker 2 (Guo et al. 2013). Processing by BS-Seeker 2 includes the removal of sequencing adapters. Methylation was called with BS-Seeker 2. DNA methylation heatmaps over genes and transposable elements were generated with the deepTools suite (Ramírez et al. 2016), by generating BIGWIG files for each sample and computing matrix with BED containing TAIR10 genic regions. Differentially methylated regions (DMRs) were identified by HOME (Srivastava et al. 2019), available on GitHub (https://github.com/Akanksha2511/HOME). DMRs were identified in CG and non-CG contexts, requiring a change in methylation levels of at least 0.1 over the whole region and an average coverage for all cytosines greater than three for both mutant and control. All annotated genes in TAIR10 were split into 3 categories based on their DNA methylation levels: genes with >2% mCHG and mCHH were classified as pseudo-genes; in the remaining list, genes with >5% mCG were labeled as GbM genes; and the rest are unmethylated genes.

#### RNA-seq

Raw sequencing data were de-multiplexed with bcl2fastq software (Illumina). FASTQ files were mapped to the TAIR10 genome using STAR with default parameters (Dobin et al. 2013). Reads underwent lightweight alignment using Salmon version 1.4.0 (Patro et al. 2017) to the TAIR10 transcriptome and reads from different transcripts originating from the same gene were combined using the R package tximport (Soneson et al. 2016) before being imported into the R package DESeq2 (Love et al. 2014) for normalisation and testing of differentially expressed genes (DEGs).

To look for novel splicing changes that occurred within the mC reader mutants, the reads mapped with STAR (Dobin et al. 2013) were processed by StringTie and merged together (Pertea et al. 2015) into a master novel transcriptome comprising splicing events from TAIR10 and ones uniquely identified within this study. Then reads underwent lightweight alignment using Salmon version 1.4.0 (Patro et al. 2017) against the novel transcriptome. Novel and known transcripts belonging to the same gene were analysed for splicing events by SUPPA2 (Trincado et al. 2018). Differential alternative splicing (DAS) was calculated for each event based on abundance of transcripts with and without inclusion of those events by SUPPA2 (Trincado et al. 2018).

#### GRO-seq

Illumina adapters were trimmed from the raw data using cutadapt (Martin 2011) with settings for trimming of sRNA libraries that were used in GRO-seq: `-a AGATCGGAAGAGCACACGTCTGAACTCCAGTCAC −m 10`. Ribosomal RNA reads were filtered from the BAM file with RSeQC python package. Reads were then mapped to the TAIR10 genome using STAR with default parameters (Dobin et al. 2013). For 5’ GRO-seq, HOMER with `-style tss` was used to define regions of transcriptional activity, meanwhile for standard GRO-seq, HOMER with `-style groseq` was used to define regions of transcriptional activity (Wang et al. 2011). Differential transcription of these transcribed regions was calculated by edgeR (Robinson et al. 2010; Nikolayeva and Robinson 2014), as done previously (Chae et al. 2015).

#### Affinity Purification-MS

We filtered out the contaminant proteins listed in (Van Leene et al. 2015), since they are common contaminating proteins from TAP-MS experiments in Arabidopsis. One experiment was performed as a single replicate (Set 1) with tandem affinity purification (TAP) with Strep and His tags, and one experiment was performed in three replicates (Set 2) with affinity purification (AP) with just the Strep tag, each followed by mass spectrometry (MS) identification of peptide fragments. After MS, the bait proteins in each experiment were the most abundant protein by peptide count (Supp Table 5), as expected. We therefore normalised the peptides of other proteins as a percentage of those identified for the bait protein (Supp Table 5), to find the most enriched proteins, which are the most likely to be true interactors. For comparison to the literature, we extracted protein-protein interactors listed in the following publications and then searched for all of these interactions that were supported by at least one peptide within either of our (T)AP-MS datasets: (Li et al. 2015, 2017; Harris et al. 2018; Zhao et al. 2019; Ichino et al. 2021a; Feng et al. 2021; Zhou et al. 2021). The rules for inclusion in our network of mC reader protein-protein interactions were:

1. If above threshold (15%) in Set1 experiment and Set2 combined replicates.
2. If above a stringent threshold (35%) in combined Set2 replicates (and over 30% in at least 2/3 individual replicates).
3. If found in other published studies as an interactor and peptides were identified in either Set1 experiment or Set2 experiment.

## Acknowledgements

J.C. was supported by a UWA International Postgraduate Research Scholarship. The Vermeulen lab is part of the Oncode Institute, which is partly funded by the Dutch Cancer Society (KWF). This work was supported by the following grants to R.L.: Australian Research Council (ARC) Centre of Excellence in Plant Energy Biology (CE140100008), ARC DP210103954, NHMRC Investigator Grant GNT1178460, Silvia and Charles Viertel Senior Medical Research Fellowship, and Howard Hughes Medical Institute International Research Scholarship.

## Authors contributions

J.C. and R.L. conceived of the project and designed the experiments. J.C. performed the ChIP-seq, DAP-seq, RNA-seq and GRO-seq experiments, J.P.1 conducted the sequencing. J.C. and O.B. performed the affinity pull-down experiment, I.D.K performed the corresponding mass-spectrometry and its analysis. J.C., J.P.B.L., P.W.T.C.J, O.D., A.H.M and J.P.2 performed the (T)AP-MS experiments and/or their analysis. J.C., J.P.B.L. and R.L. analyzed the data and/or its significance. A.H.M, M.V. and R.L. acquired funding. J.C., J.P.B.L and R.L. wrote the manuscript. All authors read and approved the final manuscript.

## List of Supplemental Materials

Supplemental Figure 1: Design of the DNA probes used in the affinity pull-down of Arabidopsis mC readers.

Supplemental Figure 2: ChIP-seq biological replicates show reproducible results.

Supplemental Figure 3: Top motifs over-represented under the peaks of each candidate.

Supplemental Figure 4: MBD2, MBD5 and MBD6 bind to gene-body methylation.

Supplemental Figure 5: The DNA methylome and transcriptome of mC reader mutant plants is highly stable.

Supplemental Figure 6: Mutations of mC readers does not impair the normal epigenome of histone modifications.

Supplemental Table 1. List of mC reader proteins enriched in binding to methylated probes and previous literature knowledge.

Supplemental Table 2. List of proteins and probe binding statistics from Figure 1b

Supplemental Table 3. List and characteristics of complemented mutant lines generated in this study.

Supplemental Table 4. Description of the T-DNA mutant lines used and CRISPR mutant lines generated for this study.

Supplemental Figure 5. List of proteins identified in each (T)AP-MS experiments.

Supplemental Table 6. List of previously published protein-protein interactions used to generate the protein network.

Supplemental Table 7. List of primers used in this study

