## Supplementary Figures for "Characterization of DNA methylation reader proteins of *Arabidopsis thaliana*"

### Supp Figures

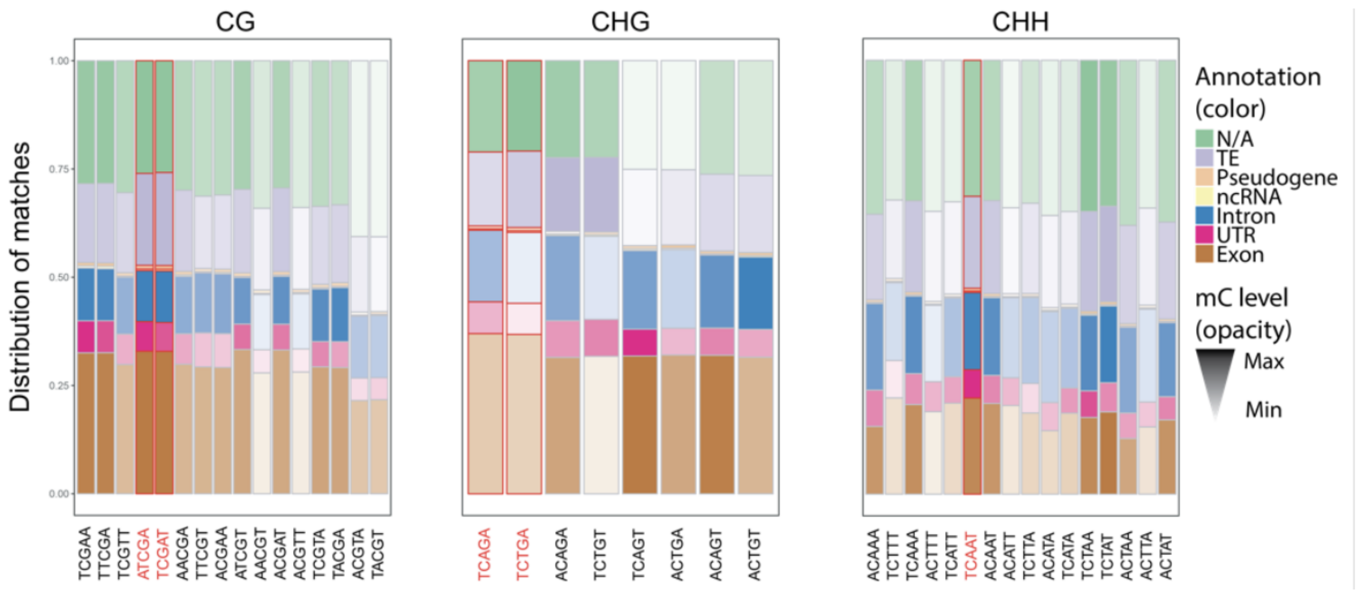


**Supplemental Figure 1: Design of the DNA probes used in the affinity pull-down of Arabidopsis mC readers.**
Distribution of matches of all the 5-bp motifs in Arabidopsis Col-0. The colour is representative of the annotated feature, the transparency is representative of the methylation level. For each feature, the motif with the highest methylation level is the darkest, the motif with the lowest methylation is the clearest. The chosen motifs are boxed and labelled in red

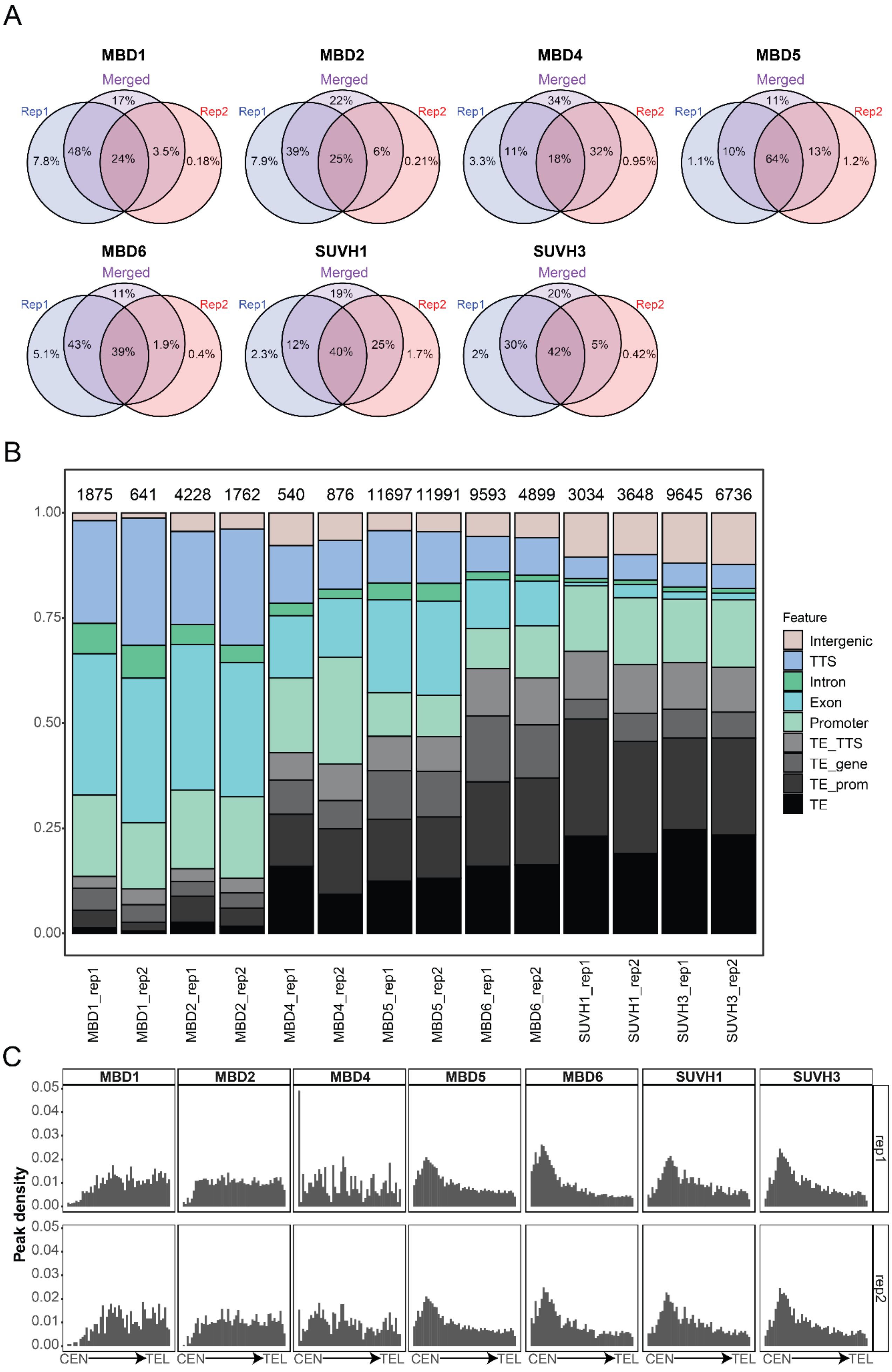
A
MBD1
Merged
Rep1
Rep2
17%
7.8%
48%
24%
3.5%
0.18%
MBD2
Merged
Rep1
Rep2
22%
7.9%
39%
25%
6%
0.21%
MBD4
Merged
Rep1
Rep2
34%
3.3%
11%
18%
32%
0.95%
MBD5
Merged
Rep1
Rep2
11%
1.1%
10%
64%
13%
1.2%
MBD6
Merged
Rep1
Rep2
11%
5.1%
43%
39%
1.9%
0.4%
SUVH1
Merged
Rep1
Rep2
19%
2.3%
12%
40%
25%
1.7%
SUVH3
Merged
Rep1
Rep2
20%
2%
30%
42%
5%
0.42%
B
1875 641 4228 1762 540 876 11697 11991 9593 4899 3034 3648 9645 6736
1.00
0.75
0.50
0.25
0.00
Feature
Intergenic
TTS
Intron
Exon
Promoter
TE_TTS
TE_gene
TE_prom
TE
MBD1_rep1
MBD1_rep2
MBD2_rep1
MBD2_rep2
MBD4_rep1
MBD4_rep2
MBD5_rep1
MBD5_rep2
MBD6_rep1
MBD6_rep2
SUVH1_rep1
SUVH1_rep2
SUVH3_rep1
SUVH3_rep2
C
MBD1
MBD2
MBD4
MBD5
MBD6
SUVH1
SUVH3
Peak density
0.05
0.04
0.03
0.02
0.01
0.00
rep1
rep2
CEN
TEL

**Supplemental Figure 2: ChIP-seq biological replicates show reproducible results.**

A. Overlap between ChIP-seq peaks called for each individual replicate and when merged before calling peaks (as percentage of the total number of individual peaks). One replicate often contributes more to the total amount due to different expression levels of the tagged-protein, but very few peaks from each replicate are lost when merging the data, showing robustness of the signal.

B. Distribution of peaks in genomic features for each biological replicate, similar to Fig 2A. Total numbers of peaks for each sample are indicated at the top.

C. Density of mC reader ChIP-seq peak position along the Arabidopsis chromosomes, from centromere (CEN) to telomere (TEL), for each biological replicate, similar to Fig 2B.

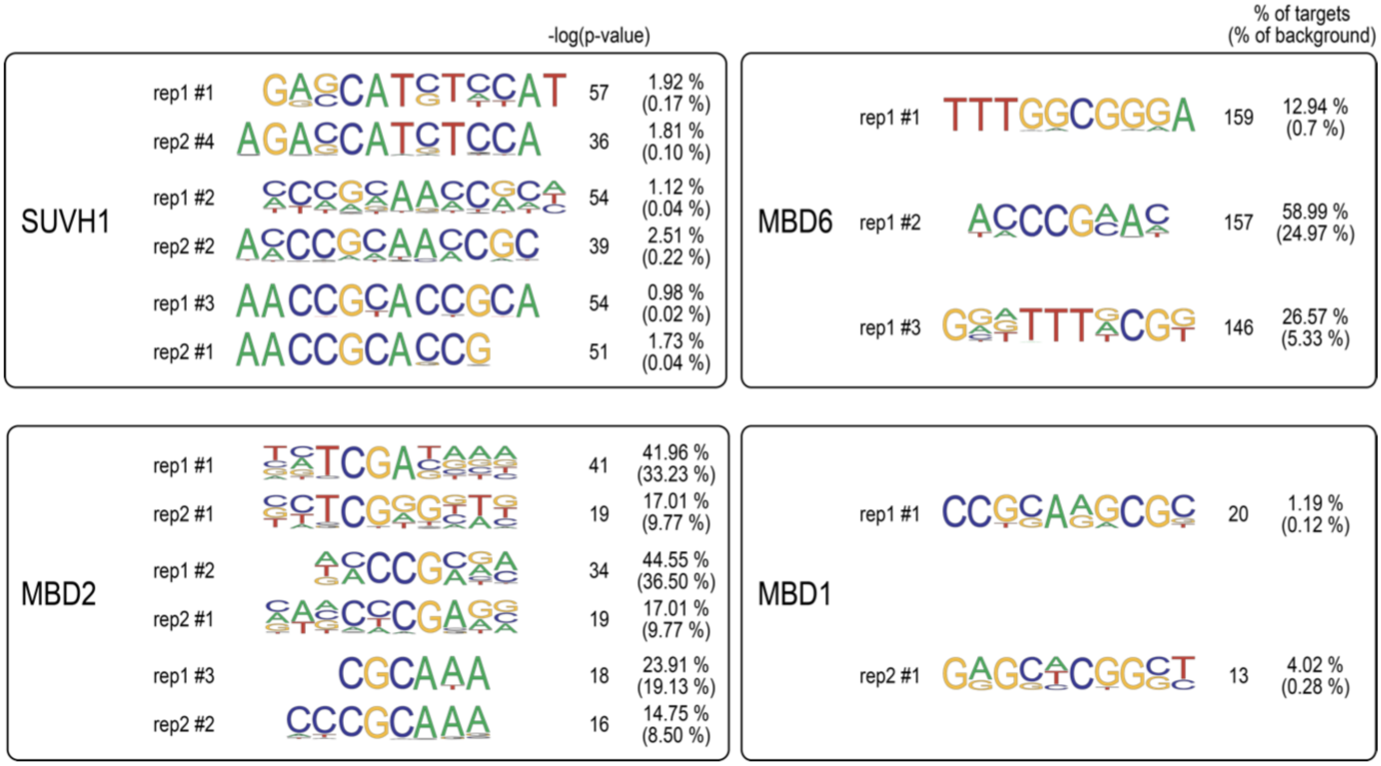


**Supplemental Figure 3: Top motifs over-represented under the peaks of each candidate.**
Peaks and motifs were identified with the Homer suite. According to Homer guidelines, only motifs with -log(p-value)>>50 can be considered real. No homology was found between MBD1 replicates, representative of a broader binding pattern, so only their respective first motifs were included.

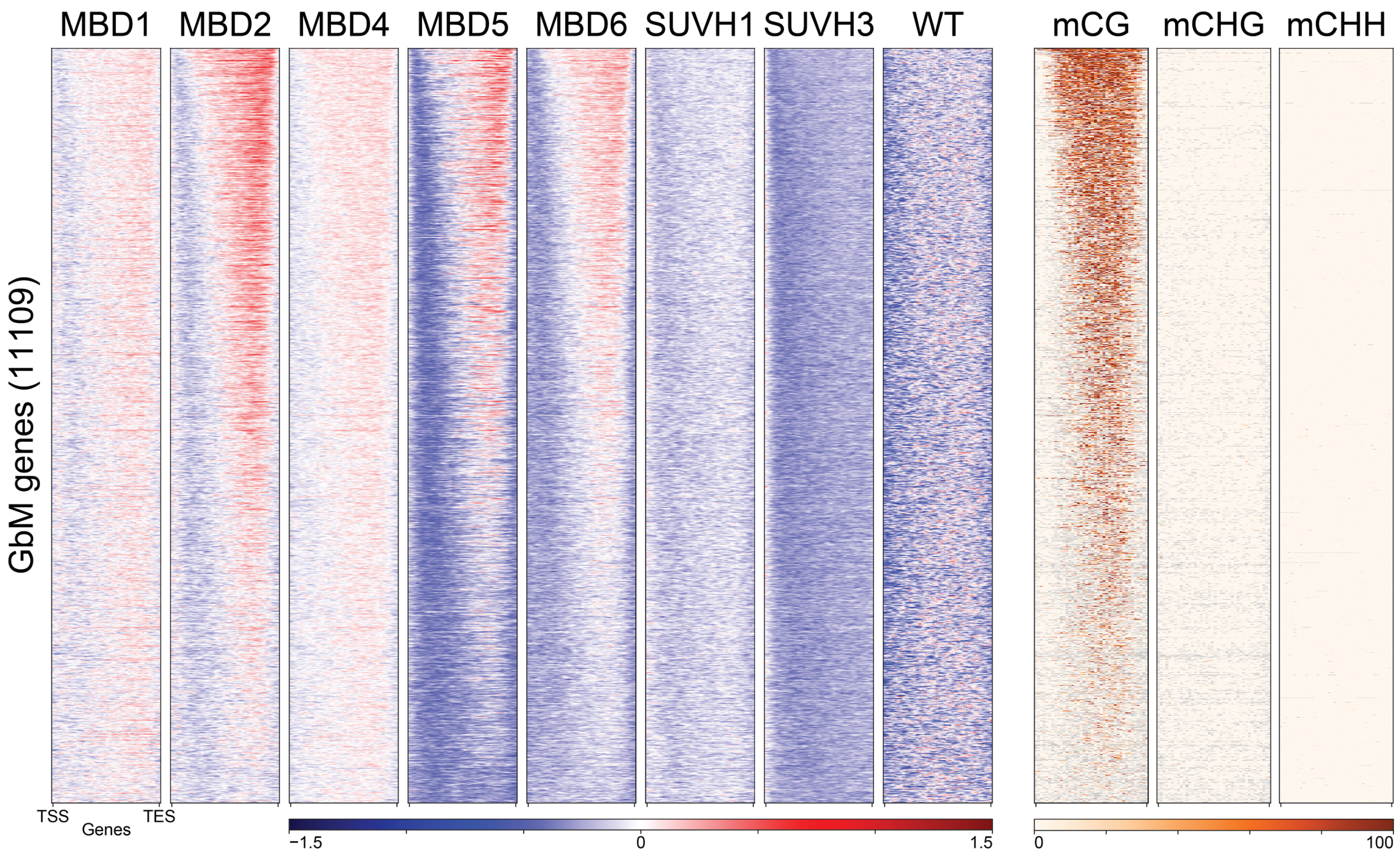


**Supplemental Figure 4: MBD2, MBD5 and MBD6 bind to gene-body methylation.**
Heatmap showing enrichment of the mC reader candidates by ChIP-seq over all annotated genes with gene-body methylation (see Methods). MBD2, MBD5 and MBD6 show enrichment at the 5' end of genes, which colocalizes with mCG levels. Heatmap generated with Deeptools, by scaling all genes to 2 kb and plotting 50 bp bins.

A

| HOME pairwise comparison | # CG DMRs |
|---|---|
| WT VS *mbd1* | 1098 |
| WT VS *mbd4* | 561 |
| WT VS *mbd6* | 1044 |
| WT VS *suvh1;3* | 1003 |
| WT VS *suvh1* | 1203 |
| WT VS *suvh3* | 1909 |
| WT VS *mbd1;2;4* | 1051 |
| WT VS *mbd1;2;5;6* | 1126 |
| WT VS *mbd2;5;6* | 1266 |
| WT VS *hdg11;12* | 1031 |
| WT VS *mbd1;2* | 875 |
| WT VS *mbd1;4* | 726 |
| WT VS *mbd2* | 1869 |
| WT VS *mbd5;6* | 815 |
| WT VS *mbd5* | 1020 |

B

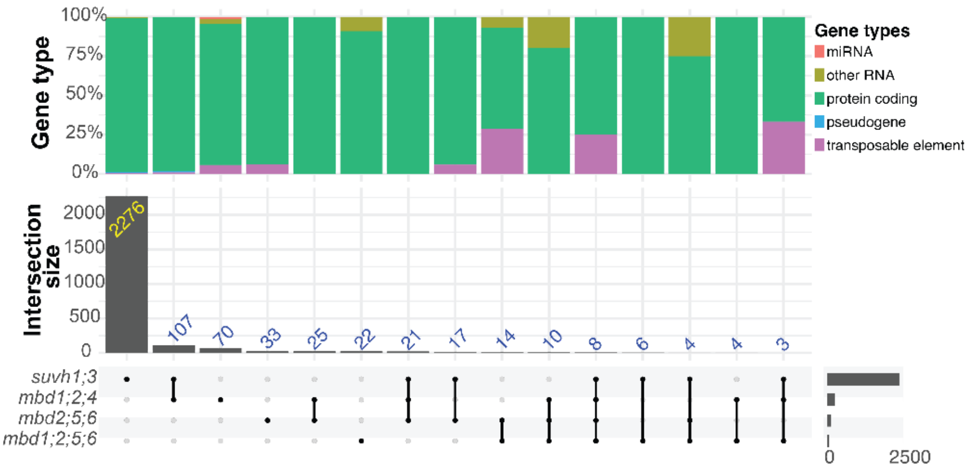


C

| DMR WT vs: | ChIP bait | CG DMRs | mC reader ChIP | Overlap |
|---|---|---|---|---|
| *mbd1;2;4* | MBD1 | 1051 | 2069 | 28 |
| | MBD2 | | 5311 | 135 |
| | MBD4 | | 1544 | 47 |
| *mbd1;2;5;6* | MBD1 | 1126 | 2069 | 39 |
| | MBD2 | | 5311 | 179 |
| | MBD5 | | 14531 | 458 |
| | MBD6 | | 10249 | 370 |
| *mbd2;5;6* | MBD2 | 1266 | 5311 | 194 |
| | MBD5 | | 14531 | 528 |
| | MBD6 | | 10249 | 412 |
| *suvh1;3* | SUVH1 | 1003 | 4913 | 153 |
| | SUVH3 | | 12164 | 242 |

D

| Sample | CG DMRs | DEG (promoter) | Overlap |
|---|---|---|---|
| *mbd1;2;4* | 1051 | 248 | 2 |
| *mbd1;2;5;6* | 1126 | 71 | 0 |
| *mbd2;5;6* | 1266 | 132 | 4 |
| *suvh1;3* | 1003 | 2442 | 24 |

E

| Sample | CG DMRs | DEG (gene body) | Overlap |
|---|---|---|---|
| *mbd1;2;4* | 1051 | 248 | 7 |
| *mbd1;2;5;6* | 1126 | 71 | 3 |
| *mbd2;5;6* | 1266 | 132 | 8 |
| *suvh1;3* | 1003 | 2442 | 38 |

F

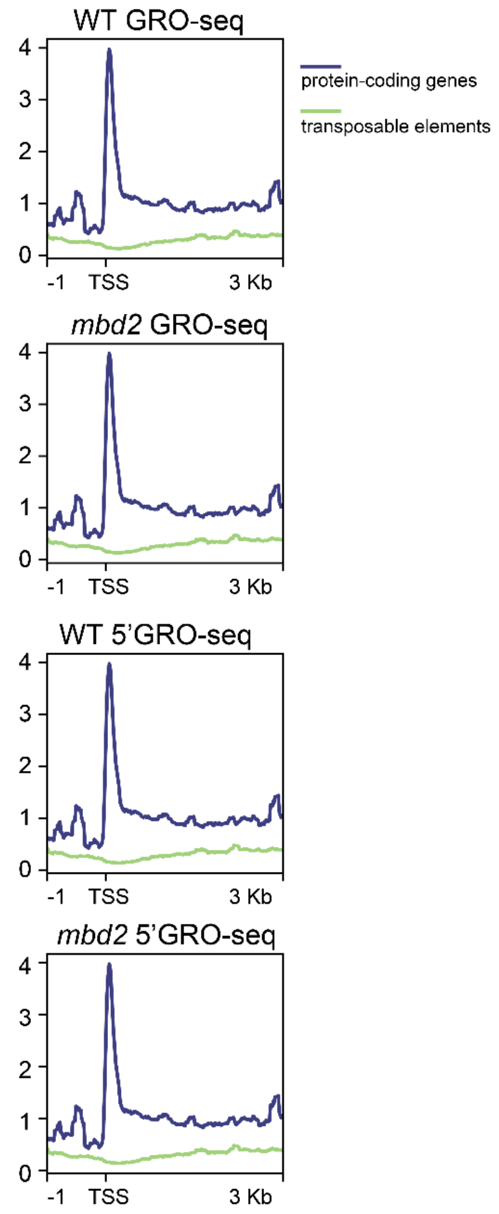


G

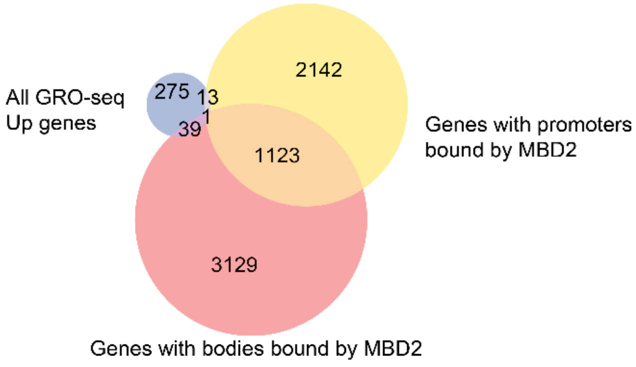

**Supplemental Figure 5: The DNA methylome and transcriptome of mC reader mutant plants is highly stable.**

A) CG-DMRs identified in the various mC reader mutant seedlings in comparison to WT seedlings.

B) The number of differentially expressed genes identified in the higher-order mutants within this study and their overlap with other mutants.

C) The overlap of CG-DMRs in higher order mutants with mC reader ChIP peaks.

D) The overlap of differentially expressed genes (DEGs) within each mutant with CG-DMRs identified within the higher order mutant that overlap the DEG’s promoter

E) The overlap of DEGs within each mutant with CG-DMRs identified within the same mutant that overlap the DEG’s gene body.

F) GRO-seq signal over TAIR10 genes, split into protein-coding genes and transposable element genes.

G) Overlapping of the genes with increased transcription in either normal GRO-seq or 5’ GRO-seq with MBD2-bound promoters or the gene bodies.

A

| | WT vs: | *mbd1* | *mbd2* | *mbd1;4* | *mbd5;6* | *mbd1;2;4* | *mbd1;2;5;6* | *mbd2;5;6* | *suvh1;3* |
|---|---|---|---|---|---|---|---|---|---|
| **Heterochromatin mark** | H3K9me2 | 7/3 | 10/8 | 22/6 | 1/1 | 35/12 | 53/8 | 26/9 | 140/35 |
| **Polycomb marks** | H2AKub | 5/3 | 2/8 | 1/0 | NA | 33/59 | 356/352 | 146/190 | 176/227 |
| | H3K27ac | 3/2 | 13/4 | 0/1 | NA | NA | NA | NA | NA |
| | H3K27me3 | 2/1 | 18/5 | 1/1 | NA | 13/24 | 52/48 | 18/55 | 21/73 |
| | H3K36me3 | 0/2 | 0/2 | 0/0 | NA | NA | NA | NA | NA |
| **TSS/enhancer marks** | H3K4me1 | 0/1 | 2/1 | 0/0 | NA | 2/0 | 4/0 | 0/0 | 0/0 |
| | H3K4me2 | 1/1 | 9/4 | 0/1 | NA | NA | NA | NA | NA |
| | H3K4me3 | 0/3 | 12/6 | 0/0 | NA | NA | NA | NA | NA |

B

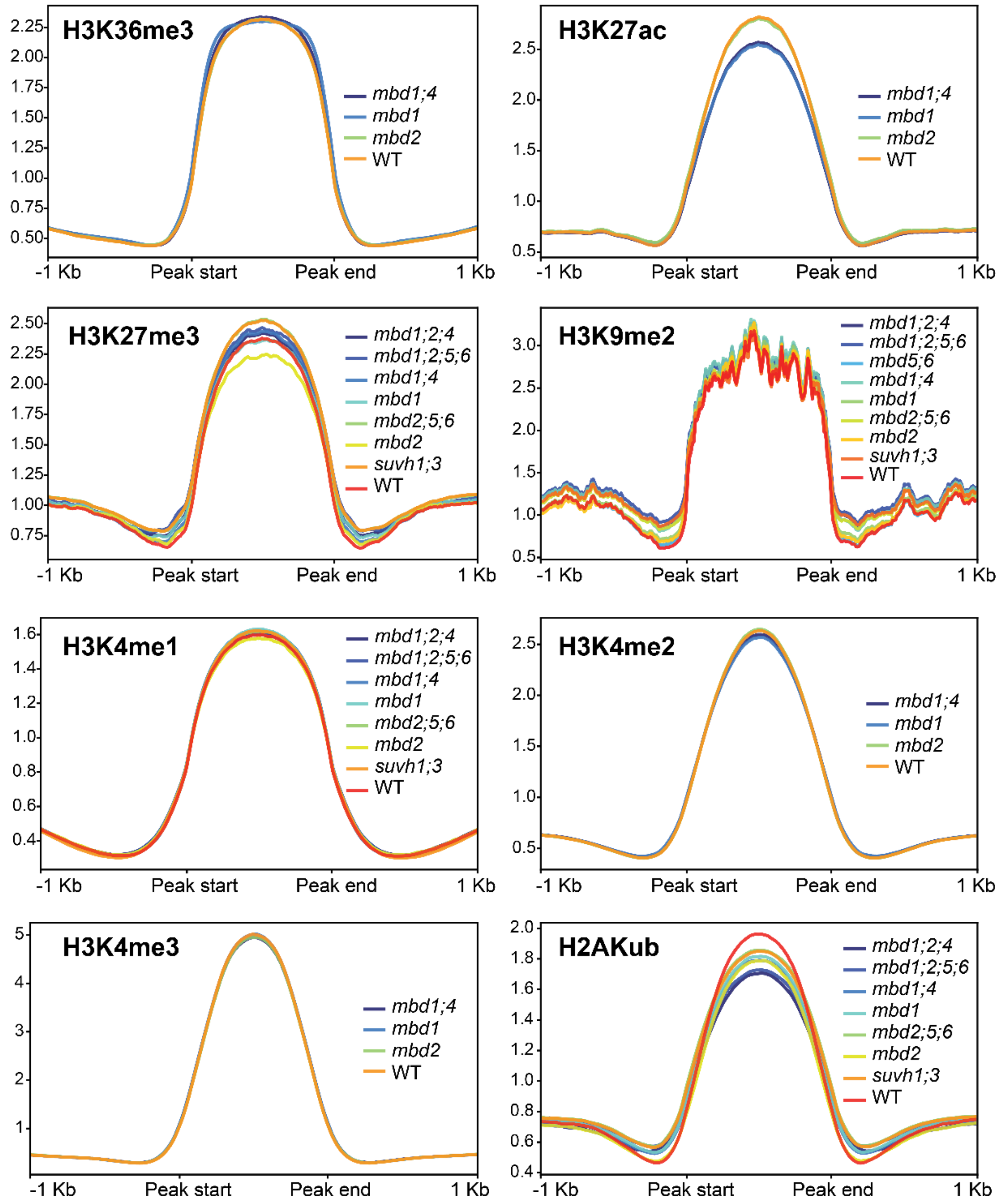


**Supplemental Figure 6: Mutations of mC readers does not impair the normal epigenome of histone modifications.**

A) Number of chromatin mark peaks that increase/decrease in each mC reader mutant. NA represents “not applicable” as that combination of mC reader mutant and histone modification/variant was not examined in this study.

B) Metaplot of histone modifications examined within this study.
